# Integrative Single-cell RNA-Seq and ATAC-Seq Analysis of Human Developmental Haematopoiesis

**DOI:** 10.1101/2020.05.06.080259

**Authors:** Anna Maria Ranzoni, Andrea Tangherloni, Ivan Berest, Simone Giovanni Riva, Brynelle Myers, Paulina M. Strzelecka, Jiarui Xu, Elisa Panada, Irina Mohorianu, Judith B. Zaugg, Ana Cvejic

## Abstract

Regulation of haematopoiesis during human development remains poorly defined. Here, we applied single-cell (sc)RNA-Seq and scATAC-Seq analysis to over 8,000 human immunophenotypic blood cells from foetal liver and bone marrow. We inferred their differentiation trajectory and identified three highly proliferative oligopotent progenitor populations downstream from haematopoietic stem cell/multipotent progenitors (HSC/MPPs). Along this trajectory, we observed opposing patterns of chromatin accessibility and differentiation that coincided with dynamic changes in the activity of distinct lineage-specific transcription factors. Integrative analysis of chromatin accessibility and gene expression revealed extensive epigenetic but not transcriptional priming of HSC/MPPs prior to their lineage commitment. Finally, we refined and functionally validated the sorting strategy for the HSC/MPPs and achieved around 90% enrichment. Our study provides a useful framework for future investigation of human developmental haematopoiesis in the context of blood pathologies and regenerative medicine.

## Introduction

During embryonic development, haematopoietic stem cells (HSCs) need to rapidly differentiate into mature blood cells. Our current knowledge of foetal haematopoietic stem and progenitor cells (HSPCs) has been mainly advanced by murine and *in vitro* model systems. It has been demonstrated that foetal haematopoiesis consists of several, separate waves of specification, migration, and differentiation of rare HSCs at distinct organs during development (Ivanovs et al., 2017). In humans, definitive haematopoiesis starts with the appearance of HSCs within haematopoietic clusters, in the dorsal aorta, at 27 days post-conception. These definitive HSCs first colonise the foetal liver at 4 post-conceptional weeks (pcw) where they expand in numbers. At 10.5 pcw, the haematopoietic site shifts once more to the cavities of bones (i.e., bone marrow), where adult haematopoiesis is established permanently. The first HSCs that seed the bone marrow are thought to continue to rapidly increase in numbers before undergoing a dramatic change in their proliferative and differentiation properties to accommodate the need for high production of differentiated progeny (Mikkola and Orkin, 2006).

Historically, differentiation processes in the haematopoietic system have been depicted as a series of intermediate steps, defined by panels of cell surface markers (i.e., cluster of differentiation, CD). In this model, often represented as a “haematopoietic tree”, HSCs give rise to increasingly lineage-restricted cell types, eventually leading to mature blood cells (Akashi et al., 1999) (Weissman, 2000). This paradigm shifted in the last five years with several studies reporting the transcriptomes of thousands of single haematopoietic cells, isolated by cell surface markers, both in the mouse model and in adult humans (Paul et al., 2015) (Velten et al., 2017). These reports showed that progenitor populations, previously thought to be homogeneous, are actually very heterogeneous on the transcriptional level.

The mechanisms underlying early fate decisions in HSCs are largely unknown. It has been postulated that the stochastic expression of lineage-specific transcription factors (TFs) above the noise threshold can “lock” a cell into a distinct cell fate (Graf and Enver, 2009). In line with this, co-expression of genes associated with antagonistic lineages, including key TFs, have been observed in multipotent haematopoietic cells, albeit at low levels (Hu et al., 1997) (Miyamoto et al., 2002). This points towards the presence of sub-populations of cells within the multipotent compartment that are permissive for opposing cell fates prior to their lineage commitment, a phenomenon referred to as priming (Nimmo et al., 2015). More recently, single-cell RNA sequencing (scRNA-Seq) of human HSPCs introduced a different concept of priming. Studies of the adult bone marrow and foetal liver haematopoiesis identified sub-populations of haematopoietic stem cells and multipotent progenitors (HSC/MPPs) with a coordinated expression of marker genes, specific for distinct uni-lineage differentiation programmes, that gradually increased along all differentiation branches (Velten et al., 2017) (Popescu et al., 2019). In addition, there are some indications that lineage priming in the HSC compartment might be happening not only on the transcriptional but also at the epigenetic level (Nimmo et al., 2015). Data from single-cell Assay for Transposase Accessible Chromatin sequencing (scATAC-Seq) of phenotypic HSPCs from the adult human bone marrow show that phenotypic multipotent progenitors have variations in chromatin accessibility consistent with a bias towards erythroid and lymphoid lineages (Buenrostro et al., 2018).

Here we performed an integrative analysis of scRNA-Seq and scATAC-Seq of more than 8,000 immunophenotypic HSPCs, from 17-22 pcw human foetal liver, femur, and hip to define transcriptional and epigenetic changes during blood differentiation. We explored lineage priming at the transcriptional and chromatin level in HSC/MPPs and refined the sorting strategy for the isolation of a highly enriched HSC/MPP population.

## Results

### Single-cell transcriptome of the haematopoietic compartment in human foetal liver and bone marrow

To capture the full repertoire of haematopoietic cells during foetal development, we single-cell sorted phenotypically defined blood populations, from matched (i.e., from the same individual) foetal livers, femur, and hip (iliac) bones, between 17 and 22 pcw (Figure 1A). Cells from the liver, hip, and femur were sorted and processed independently in all experiments. Thus, each cell can be traced back to the foetus and organ it came from. We used a hierarchical approach, where we first isolated non-committed (Lin-[CD3, CD8, CD11b, CD14, CD19, and CD56] CD34+ CD38−) progenitors, that contain all immature haematopoietic populations and are present at the frequency of less than 0.1% of the total foetal bone marrow (Golfier et al., 2008), followed by a more restrictive panel to capture differentiated and mature cell types. We next isolated committed (Lin-, CD34+ CD38+) progenitors as well as phenotypic HSCs, multipotent progenitors (MPPs), common myeloid progenitors (CMPs), megakaryocyte-erythroid progenitors (MEPs), granulocyte-monocyte progenitors (GMPs), and common lymphoid progenitors (CLPs). In addition, based on broad phenotypic markers, we sorted T cells, NK cells, innate lymphoid cells (ILCs), monocytes, dendritic cells, mast cells, basophils, neutrophils, eosinophils, erythroid progenitors, erythrocytes, immature megakaryocytes (MKs), mature MKs, progenitor B cells (pro-B cells), precursor B cells (pre-B cells), mature B cells, and endothelial cells (Supplementary table 1, Supplementary figure 1).

**Figure 1.**
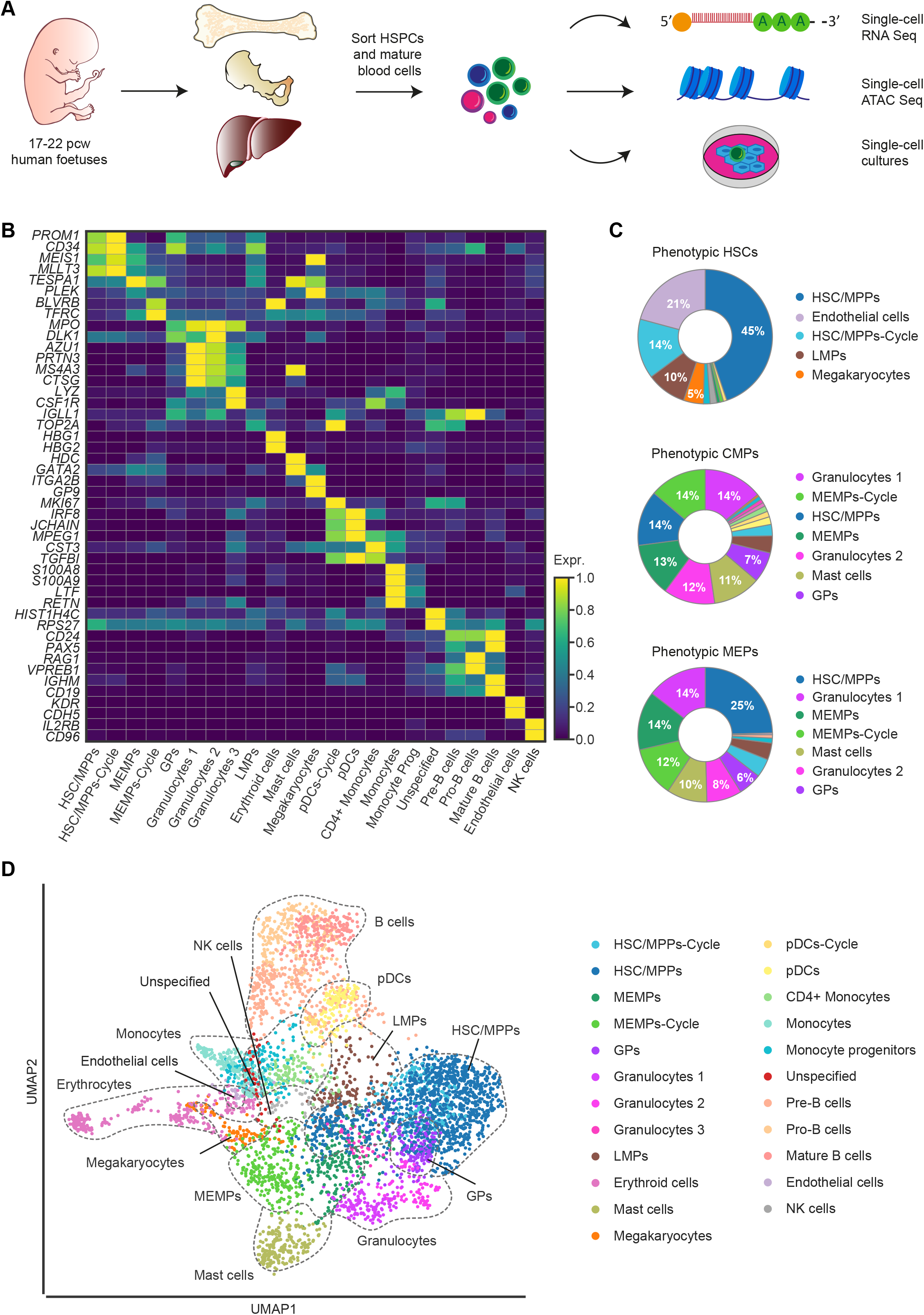
Single-cell transcriptome analysis of human foetal haematopoiesis. **A.**Schematic overview of the experimental workflow. From each foetus (age 17-22 pcw), phenotypically defined HSPCs and mature blood cells were sorted from bone marrow (femur and hip) and liver and processed for scRNA-Seq (*n*=15), scATAC-Seq (*n*=3), as well as for single-cell *in vitro* differentiation assays (*n*=4). **B.** Heatmap of the mean expression value of two manually selected marker genes for each cell type. The expression of the genes is standardised between 0 and 1. For each gene, the minimum value is subtracted and the result is divided by the maximum. The standardised expression level is indicated by colour intensity. **C.** Donut plots showing the percentage of transcriptionally defined (i.e., manually curated) cell populations in each of the phenotypically defined stem and progenitor populations. The colours correspond to the identified cell types. **D.** UMAP visualisation of haematopoietic cells from liver and bone marrow coloured by cell type. HSC/MPPs-Cycle - cycling haematopoietic stem cells/multipotent progenitors; HSC/MPPs - haematopoietic stem cells/multipotent progenitors; MEMPs - megakaryocyte-erythroid-mast progenitors; MEMPs-Cycle - cycling megakaryocyte-erythroid-mast progenitors; GPs - granulocytic progenitors; LMPs - lympho-myeloid progenitors; pDCs-Cycle - cycling plasmacytoid dendritic cells; pDCs - plasmacytoid dendritic cells.

Single cells from 15 foetuses were processed for scRNA-Seq using the SmartSeq2 protocol (Picelli et al., 2014) (Figure 1A). Overall, 4,504 cells passed quality control (QC) (Supplementary table 2) with an average of ~3,600 genes per cell and ~670,000 reads per cell (Supplementary figure 2A-C, K-L). To exclude technical batch effects, we merged the datasets from all samples and tissues using autoencoders (AEs) and applied the batch balanced *k* nearest neighbours (BBKNN) approach (Polański et al., 2020; Luecken et al., 2020) to the latent space (Tangherloni et al., 2019) (Supplementary figure 2O). We applied the graph-based Leiden clustering algorithm (Traag et al., 2019) to the batch corrected neighbourhood graph. Based on differential expression (DE) analysis and top 20 marker genes (Figure 1B) ranked on the significance of standardised expression, we manually annotated 23 distinct populations. Within the haematopoietic progenitor compartment, we annotated clusters as HSC/MPPs, HSC/MPPs-Cycle, lympho-myeloid progenitors (LMPs), megakaryocyte-erythroid-mast progenitors (MEMPs), MEMPs-Cycle, granulocytic progenitors (GPs), as well as numerous mature blood cell types as shown in the Uniform Manifold Approximation and Projection (UMAP) space (Becht et al., 2018) (Figure 1D).

Of the mature blood cell types, we identified clear transcriptional signatures of erythroid cells (expressing *HBG1*, *HBA1*, *GYPA*, and *ALAS2*), megakaryocytes (expressing *FLI1*, *ITGA2B*, and *GP9*), monocyte progenitors and monocytes (expressing *CD14*, *MPEG1*, and *CD33*), CD4+ monocytes, mast cells (expressing *CD63*, *GATA2*, and *HDC*), plasmacytoid dendritic cells (pDCs - expressing *IL3RA*, *IRF8*, *MPEG1*, and *JCHAIN*), with an additional cluster of highly cycling pDCs (expressing pDC and proliferation markers, e.g., *MKI67*) and granulocytes 1, 2, and 3 (expressing *AZU1*, *MPO*, and *PRTN3*), (Figure 1B, Supplementary figure 3). While granulocytes were present in our dataset, we could not clearly distinguish neutrophils, basophils, and eosinophils due to the mixed expression signatures. In the lymphoid compartment, we identified NK cells (expressing *CD3D*, *IL2RB*, and *CD96*) and B cells (expressing *CD19* and *CD79B*) (Figure 1B, Supplementary figure 3). B cell lineage included pro-B, which showed expression of *IGLL1* and *RAG1*, and pre-B, expressing high levels of *CD79B*, *VPREB1*, and *CD24* (Figure 1B). Finally, we identified a cluster of mature B cells, expressing high levels of *IGHM* and decreased levels of *IGLL1*, compared to pro/pre B clusters (Figure 1B). We did not detect any T cells or ILCs in the liver or in the femur in spite of sorting phenotypic T cells and ILCs using broad cell surface markers for these populations. Unlike B cells that mature in the BM, T cells derive from lymphoid progenitors that migrate from the BM to the thymus, where they complete their maturation. The development of ILCs is less understood but there have been suggestions that ILC precursors migrate early on from BM into non-haematopoietic tissues, e.g., gut (Cichocki et al., 2019). Since we only sorted BM and not thymus or gut, we might have captured only progenitors but not T cells and ILCs. By using a Deep Neural Network (DNN) (LeCun et al., 2015) and the top 30 marker genes for each cluster, we were able to correctly classify the cells to the prospective clusters with 90.46% accuracy, confirming that our manual annotation of clusters separated well the distinct cell types/states (see Methods, Supplementary figure 4A).

In the last decade, human HSCs and other progenitor populations have been isolated and used in functional assays based on specific sets of cell surface markers. It has been suggested that the foetal haematopoietic progenitor compartment differs substantially from its adult counterpart (Notta et al., 2016). Our approach allowed us to compare the extent to which the phenotypic identity of cell populations (as defined by CD markers) matched their transcriptional state, i.e., our manually curated clusters and thus to critically examine the use of CD markers in the context of foetal bone marrow haematopoiesis.

Single-cell analysis revealed substantial transcriptional heterogeneity within all immunophenotypically-defined stem and progenitor populations, with some phenotypic progenitor populations such as HSCs, MPPs, CMPs, GMPs, MEPs, and CLPs being comprised of more than ten different transcriptionally-defined populations. (Figure 1C, Supplementary figure 4B). This observation is in agreement with recent research showing a high level of heterogeneity of the progenitor compartment of human cord blood (Knapp et al., 2018). Taken together, our comparative analysis shows that currently used cell-surface markers are a poor predictor of the transcriptional state of human foetal haematopoietic progenitors.

### Inference of differentiation trajectories during foetal haematopoiesis

Next, we used a Force-Directed Graph drawing algorithm, ForceAtlas2, to infer the differentiation trajectory of haematopoietic cells during human foetal development (Jacomy et al., 2014). We initialised a ForceAtlas2 layout with Partition-based Approximate Graph Abstraction (PAGA) coordinates from our annotated cell types (Wolf et al., 2019). This initialisation generated an interpretable single-cell embedding that is faithful to the global topology. The obtained global topology revealed HSC/MPPs at the tip of the trajectory (Figure 2A-B, Supplementary figure 3). HSC/MPPs showed high expression of *MLLT3*, a crucial regulator of human HSC maintenance (Calvanese et al., 2019), *HLF*, a TF involved in preserving quiescence in HSCs (Komorowska et al., 2017) and *MEIS1*, a TF involved in limiting oxidative stress in HSCs, which is necessary for quiescence (Unnisa et al., 2012) (Wang et al., 2018). Cells in this cluster also expressed high levels of surface markers of HSPCs such as *CD34*, (Morisot et al., 2006), *SELL* (Ivanovs et al., 2017), and *PROM1* (de Wynter et al., 1998) (Saha et al., 2020) (Figure 1B and 2C). Downstream of HSC/MPPs, we identified three distinct, highly proliferative, oligopotent progenitor populations. We used Scanpy's *dpt* function to infer the progression of the cells through geodesic distance along the graph. Then, we used Scanpy's *paga_path* function to show how the gene expression and annotation changes along the three main paths (MEMPs, GP, and LMPs) that are present in the abstracted graph (Figure 2C).

**Figure 2.**
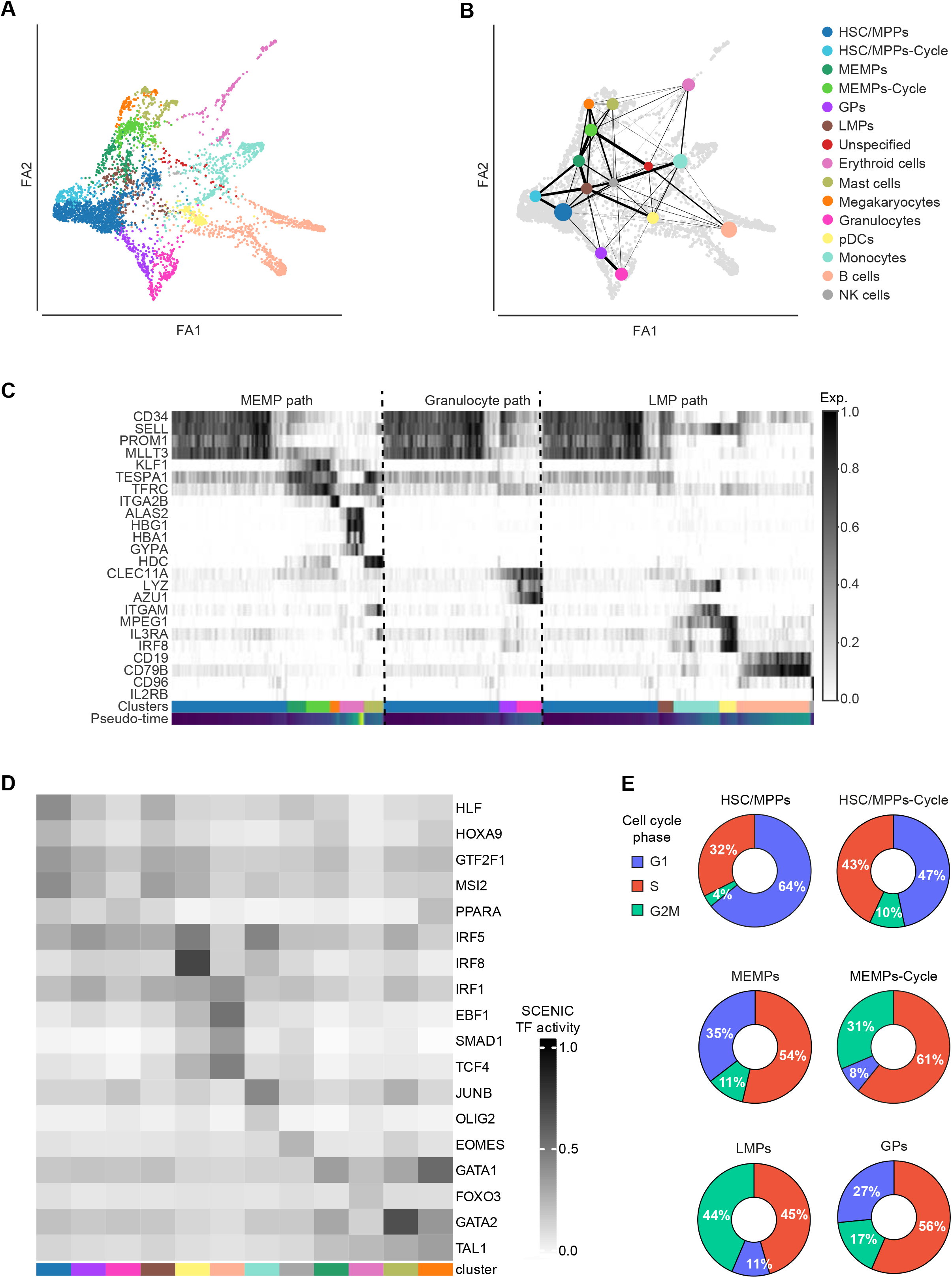
Differentiation trajectory of human foetal haematopoietic cells. **A.** FDG visualisation of the differentiation trajectory of haematopoietic cells from Figure 1D. **B.** PAGA trajectory model imposed on the FDG visualisation of the differentiation trajectory. The size of the dots is proportional to the number of cells in the clusters. **C.** Heatmap showing dynamic expression of lineage-specific genes along the three differentiation paths (MEMP, Granulocyte, and LMP path). Cluster colours match those of **(A)** and **(B)**. **D.** Heatmap of the normalised AUC score of selected TFs for each cell type obtained by pySCENIC. Cluster colours match those of **(A)** and **(B)**. **E.** Donut plots showing the percentages of cells in G1, S, and G2M in HSC/MPPs, HSC/MPPs-Cycle, MEMPs, MEMPs-Cycle, LMPs, and GPs. Force-Directed Graph - FDG; ForceAtlas2 - FA2; HSC/MPPs-Cycle - cycling haematopoietic stem cells/multipotent progenitors; HSC/MPPs - haematopoietic stem cells/multipotent progenitors; MEMPs - megakaryocyte-erythroid-mast progenitors; MEMPs-Cycle - cycling megakaryocyte-erythroid-mast progenitors; GPs - granulocytic progenitors; LMPs - lympho-myeloid progenitors; pDCs-Cycle - cycling plasmacytoid dendritic cells; pDCs - plasmacytoid dendritic cells.

MEMPs connected HSC/MPPs with megakaryocytes, erythroid, and mast cells. In line with this, differentially regulated genes in the HSC/MPPs transition to MEMPs included megakaryocyte/erythroid/mast cells lineage-specific genes such as *GATA1*, *ITGA2B, PLEK*, *KLF1*, *HDC, and MS4A3*, (Figure 1B, 2C, Supplementary figure 5B). Presence of MEMPs in our dataset is consistent with studies in mouse models proposing a common trajectory between erythroid, megakaryocytic, and mast cell lineages (Franco et al., 2010). This concept was more recently supported by a study in human foetal liver showing a shared progenitor of megakaryocyte, erythroid, and mast cells (Popescu et al., 2019). In addition, we identified a proliferative population of MEMPs-Cycle of which ~92% were in the G2M/S phase compared to 65% of MEMPs (Figure 2E). MEMPs-Cycle population further upregulated erythroid-specific genes such as *KLF1*, *BLVRB*, and *TFRC* compared to MEMPs suggesting their gradual commitment towards erythroid lineage (Supplementary figure 5C).

GPs connected the HSC/MPP cluster with granulocyte clusters. Cells in this cluster differentially expressed myeloid lineage-specific genes (e.g., *AZU1*, *LYZ*, and *MPO*) compared to HSC/MMPs (Figure 2C, Supplementary figure 5D and were highly cycling, with 73% of cells in the G2M/S phase (Figure 2E). Finally, our data pointed towards the existence of a common progenitor population for B cells, monocytes, pDCs, and NK cells, here annotated as lymphoid-myeloid progenitors (LMPs). Cells in this cluster expressed genes specific to those lineages, including *IGLL1*, *HMGB2,* and *CD79B* (lymphoid) (Figure 2C, Supplementary figure 3) and upregulated lymphoid genes such as *CD81*, *IGLL1,* and *HMGN2* compared to HSC/MPP cluster (Supplementary figure 5E). Again, this was a highly proliferative population of cells with ~89% of cells being in the G2M/S phase (Figure 2E).

Our findings support previous studies on early lymphoid commitment in human cord blood, both *in vitro* and *in vivo*, which identified a shared lineage progenitor between lymphoid, NK, B, and T cells, monocytes, and dendritic cells (Doulatov et al., 2010) (Collin et al., 2011). Interestingly, the LMP cluster had higher expression of MPP-related genes such as *SPINK2*, *CD52,* and *SELL* compared to MEMPs, suggesting that these progenitors represent a more immature population compared to MEMPs (Supplementary figure 5F).

Next, we used the Python implementation of Single-Cell rEgulatory Network Inference and Clustering (SCENIC) (Aibar et al., 2017, Van de Sande et al., 2020) to identify master regulators and gene regulatory networks (GRN) in HSPC and mature blood cells across differentiation trajectories. We found 162 regulons of which some were enriched across many different cell types, often as a part of the particular differentiation branch, and some were cell-type specific (Figure 2D). We identified HLF and HOXA9 as main regulons in HSC/MPPs, whereas GATA1, GATA2, and TAL1 were identified in the MEMP branch of the haematopoietic tree (Figure 2D). FOXO3 was highly specific for erythroid cells whereas EOMES, OLIG2, and IRF8 for NK cells, monocytes, and pDCs, respectively. Importantly, the regulons confirmed the inferred differentiation trajectory.

To further explore heterogeneity within the HSC/MPP population we examined whether HSC/MPP cells simultaneously primed several different lineage-affiliated programs of gene activity. While HSC/MPPs sporadically expressed lymphoid, myeloid, or megakaryocyte-erythroid differentiation genes, we did not observe consistent expression of antagonistic lineage-affiliated genes in individual cells. In addition, after further sub-clustering the HSC/MPPs, there was no evident consolidation of lineage-affiliated transcriptional programs in any of the sub-populations (Supplementary figure 6). Our scRNA-Seq data, thus, do not support recently reported transcriptional lineage priming in the foetal HSC/MPP compartment (Popescu et al., 2019) and suggest that, transcriptionally, our HSC/MPP cluster represents a highly immature population of cells.

DE analysis between HSC/MPPs-Cycle and HSC/MPPs revealed upregulation of genes involved in cell cycle regulation (*FOS*, *PTP4A1*, *MCL1,* and *PKN2*) in HSC/MPPs-Cycle (Supplementary figure 5A) thus confirming that they are indeed a population of cycling stem and multipotent cells. In line with this, cell cycle analysis confirmed that ~36% of HSC/MPPs were cycling compared to ~53% of HSC/MPPs-Cycle (Figure 2E). The HSC/MPPs-Cycle had an increased expression of genes involved in glycolysis, a feature commonly found in proliferating cells (Ito and Suda, 2014) (Supplementary figure 5G). However, there were no other transcriptional differences between HSC/MPPs and HSC/MPP-Cycle, excluding the presence of transcriptional priming in the HSC/MPP-Cycle cluster.

Previous research showed that, contrary to adult blood progenitors that are mainly unilineage, foetal liver blood progenitors maintain multilineage potential (Notta et al., 2016). Our data are consistent with this observation and point towards the existence of three oligopotent progenitor populations downstream of the HSC/MPP compartments: MEMPs giving rise to erythroid, megakaryocytes, and mast cells, GPs differentiating into granulocytes and LMPs generating lymphoid, monocytes and dendritic cells.

### scATAC-Seq of foetal non-committed progenitors (CD34+ CD38−)

Detection of low abundant transcripts, such as TFs, might be difficult in scRNA-Seq data due to technical limitations of the approach, leading to false negatives (so-called drop-outs). The activity of these TFs can be inferred, however, from chromatin accessibility, emphasizing the importance of approaches integrating scRNA-Seq and scATAC-Seq data. In addition, chromatin accessibility at regulatory regions might precede gene activity and thus have predictive value for future transcription of a gene. Therefore, to further investigate the regulatory events in the very immature cell populations, we examined the single-cell chromatin accessibility landscape (using scATAC-Seq) of human foetal Lin- CD34+ CD38− cells (see Methods). We sequenced 4,001 cells from the liver and femur of three foetuses, 18, 20, and 21 pcw (see Methods). Based on our scRNA-Seq data, we expected that 90% of captured cells would be associated with one of the six populations: HSC/MPPs, HSC/MPPs-Cycle, MEMPs, MEMPs-Cycle, GPs, and LMPs, with HSC/MPPs(Cycle) constituting the majority (Supplementary figure 4B).

To capture peaks that are present in less abundant cell types such as MEMPs, MEMPs-Cycle, GPs, and LMPs, we employed an iterative peak-calling approach. We first defined open chromatin regions by pooling all the data and calling peaks in the pooled samples. Following dimensionality reduction with Diffusion maps (Haghverdi et al., 2015) and clustering using the Louvain community detection algorithm (Blondel et al., 2008), we performed a second round of peak calling in the clusters with more than 50 cells. Out of the initial ~474,000 reads, after preprocessing steps, (Supplementary figure 2D-F), on average we detected ~32,400 fragments per cell and 56% of those mapped to peaks (Supplementary figure 2G-H, M). Following filtering steps (Supplementary figure 2I-J, N), 3,611 cells passed QC with 152,282 distinct peaks.

### Motif accessibility dynamics along the inferred differentiation trajectories

In order to merge samples and remove the batch effects, we applied Harmony (Korsunsky et al., 2019; Luecken et al., 2020) on the first 50 Latent Semantic Indexing (LSI) components, excluding the first one because it was highly correlated to the sequencing depth (Supplementary figure 2P-Q). By using a shared nearest neighbour (SNN) modularity optimization based clustering algorithm, we obtained seven distinct clusters of differentially accessible peaks (Figure 3A).

**Figure 3.**
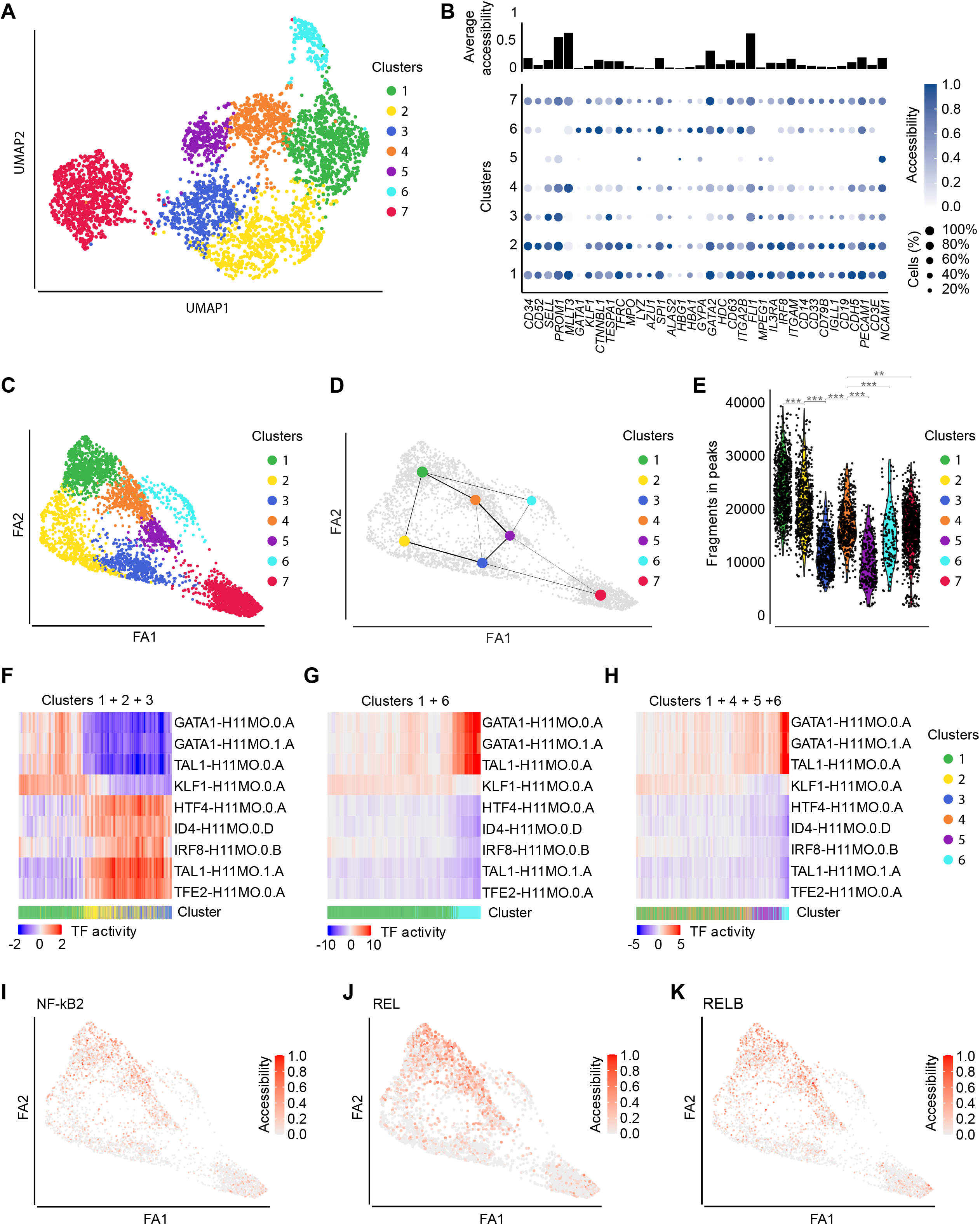
Single-cell chromatin accessibility analysis of human foetal haematopoiesis. **A.** UMAP visualisation of the scATAC-Seq dataset (=3,611 nuclei from CD34+ CD38− cells from the liver and bone marrow) coloured by cluster. **B.** (Top) Bar plot showing the average accessibility of 36 selected marker genes from our scRNA-Seq data considering all cells. (Bottom) Dot plot of the standardised accessibility of the marker genes (gene body ± 3 kb) in each of the seven clusters. For each gene, the minimum value of its accessibility is subtracted and the result is divided by the maximum value of its accessibility. The dot size indicates the percentage of cells in each cluster in which the gene of interest is accessible. **C.** FDG visualisation of the differentiation trajectory of haematopoietic cells from **(A)**. **D.** PAGA trajectory model imposed on the FDG visualisation of the differentiation trajectory of haematopoietic cells from **(A)**. The size of the dots is proportional to the number of cells in the clusters. **E.** Violin plots showing the chromatin accessibility in different clusters. p-value_1,2_<2×10^-16^; p-value_2,3_<2×10^-16^; p-value_3,4_<2×10^-16^; p-value_4,5_ <2×10^-16^; p-value_4,6_= 1.4E-07; p-value_4,7_= 0.00235. p-value<0.001 (***); 0.001<p-value<0.01 (**); 0.01<p-value<0.05 (*); p-value≥0.05 (ns). **F-H.** Heatmap showing the activity of lineage-specific TFs along differentiation trajectories: **F.** Clusters 1-2-3. **G.** Clusters 1-6. **H.** Clusters 1-4-5-6. **(I-K)** FDG visualisation of the min-max normalised TF motif accessibility along the differentiation trajectory. **I.** NF-kB2. **J.** REL. **K.** RELB. Force-Directed Graph - FDG; ForceAtlas2 - FA2.

To explore the chromatin accessibility profiles across the seven clusters, we examined the accessibility of selected marker genes from our scRNA-Seq data (Figure 3B). We observed higher accessibility of marker genes associated with stem cells (e.g., *MLLT3*, *PROM1*, *FLI1*, and *GATA2*) and lower accessibility of genes associated with distinct lineages (e.g., *MPO*, *ALAS2*, *MPEG1*, and *CD19*), keeping in line with the undifferentiated nature of sorted cells (Figure 3B). Interestingly, we observed a clear separation of clusters in terms of their overall accessibility of marker genes, with clusters 1, 2, 4, and 7 being more accessible and clusters 3 and 5 being less accessible. Cluster 6 had a mixed signature (Figure 3B).

Extensively open chromatin in multipotent cells has been previously associated with a permissive state to which multiple programmes of gene regulation may be applied upon differentiation and is considered important for the maintenance of pluripotency (Gaspar-Maia et al., 2011). To further investigate if there were global dynamic changes in accessibility patterns associated with the differentiation of foetal HSC/MPPs, we inferred differentiation pseudotime from our scATAC-Seq data using the same approach as with scRNA-Seq described above. Briefly, we built a Force-directed Graph from our seven scATAC-Seq clusters by initialising a ForceAtlas2 layout with PAGA coordinates (Figure 3C-D). The generated trajectory revealed two branches with a clear trend between chromatin accessibility and differentiation in each branch (Figure 3D-E). We observed the highest accessibility in clusters 1, 2, and 4 that gradually decreased towards the tips of the two branches (i.e., clusters 1-2 3 on one side, and 1-4-5-6 and 1-6 on the other), (Figure 3E). This result is compatible with the notion that clusters 1, 2, and 4 represent HSC/MPP population.

Control of gene expression is a dynamic process that involves both the cell-type-specific expression of TFs and the establishment of an accessible chromatin state that permits binding of TFs to a defined motif. Thus, to assess regulatory programs that are active in HSPCs, we used chromVAR (Schep et al., 2017) to calculate the most variable accessible TF sequence motifs in different clusters and examine their activity along the differentiation trajectory. Along the two branches identified by the trajectory inference, we observed dynamic changes in the accessibility of lineage-specific haematopoietic TF motifs such as GATA1, TAL1, KLF1, HTF4, ID4, IRF8, TFE2 (Figure 3F-H).

GATA1 activity (Figure 3G-H) and gene-body accessibility (Figure 3B) were enriched in cluster 6. GATA1 is known to be an important regulator of erythroid, megakaryocytic, and mast cells differentiation (Katsumura et al., 2017) and was exclusively expressed in the MEMP cluster, in our scRNA-Seq dataset. Thus, the identified trajectories between clusters 1-6 and clusters 1-4-5-6 most likely represent the MEMP differentiation paths (Figure 3D). Interestingly, in cluster 6, compared to clusters 2 and 3, we detected opposing patterns of motif accessibility for the two different TAL1 binding sites (TAL1.0.A and TAL1.1.A, respectively) (Figure 3F-H). Substantial changes in occupancy by TAL1 during differentiation have been observed, which are dependent on its binding partners (Wu et al., 2014). It has been previously reported that TAL1.0.A was co-occupied by TAL1 and GATA1 (Kassouf et al., 2010) whereas TAL1.1.A by TAL1 and TCF3 (Hsu et al., 1994). Our analysis, thus, revealed that the two different TAL1 binding motifs are active in distinct haematopoietic progenitor populations during foetal haematopoiesis (Figure 3F-H).

Clusters 2 and 3 also showed increased activity of CEBPD and IRF8, crucial for myeloid and dendritic cell differentiation and of ID4 and HTF4, which are involved in the establishment of the lymphoid lineage (Miyazaki et al., 2017) (Figure 3F-H). This points towards clusters 1, 2 and 3 forming a common initial trajectory between the myeloid and lymphoid fate, consistent with our observations in scRNA-Seq data. Cluster 1 and 4 were characterised by a high level of activity of TFs of the NF-kB pathway (i.e., NF-kB2, REL, and RELB), (Figure 3I-K), known to be involved in the regulation of HSCs maintenance and self-renewal (Zhao et al., 2012) (Espín-Palazón and Traver, 2016).

### Integrating scRNA-Seq and scATAC-Seq data

Next, we wanted to map the cells from our scATAC-Seq data to specific cell types. Since, currently, no chromatin accessibility maps are available for human foetal HSPCs, we chose a strategy to integrate our scRNA-Seq and scATAC-Seq by mapping cells based on their gene body accessibility. We used a recently developed method which identifies pairwise correspondences (termed “anchors”) between single cells across two different types of datasets, and their transformation into the shared space (Stuart et al., 2019). This approach allowed us to transfer scRNA-Seq derived annotations, learned by a classifier, onto scATAC-Seq data (see Methods).

We trained the classifier on CD34+ CD38− cells from the scRNA-Seq experiment using the six most abundant cell types (see Methods). Overall, ~57% of scATAC-Seq cells were assigned to the HSC/MPP cluster, ~18% to HSC/MPPs-Cycle, ~5% to MEMPs, ~7% to MEMPs-Cycle, ~7% to GPs, and ~3% to LMPs. Cells with the prediction score lower than 40% were labelled as unclassified (~5%) (Figure 4A).

**Figure 4.**
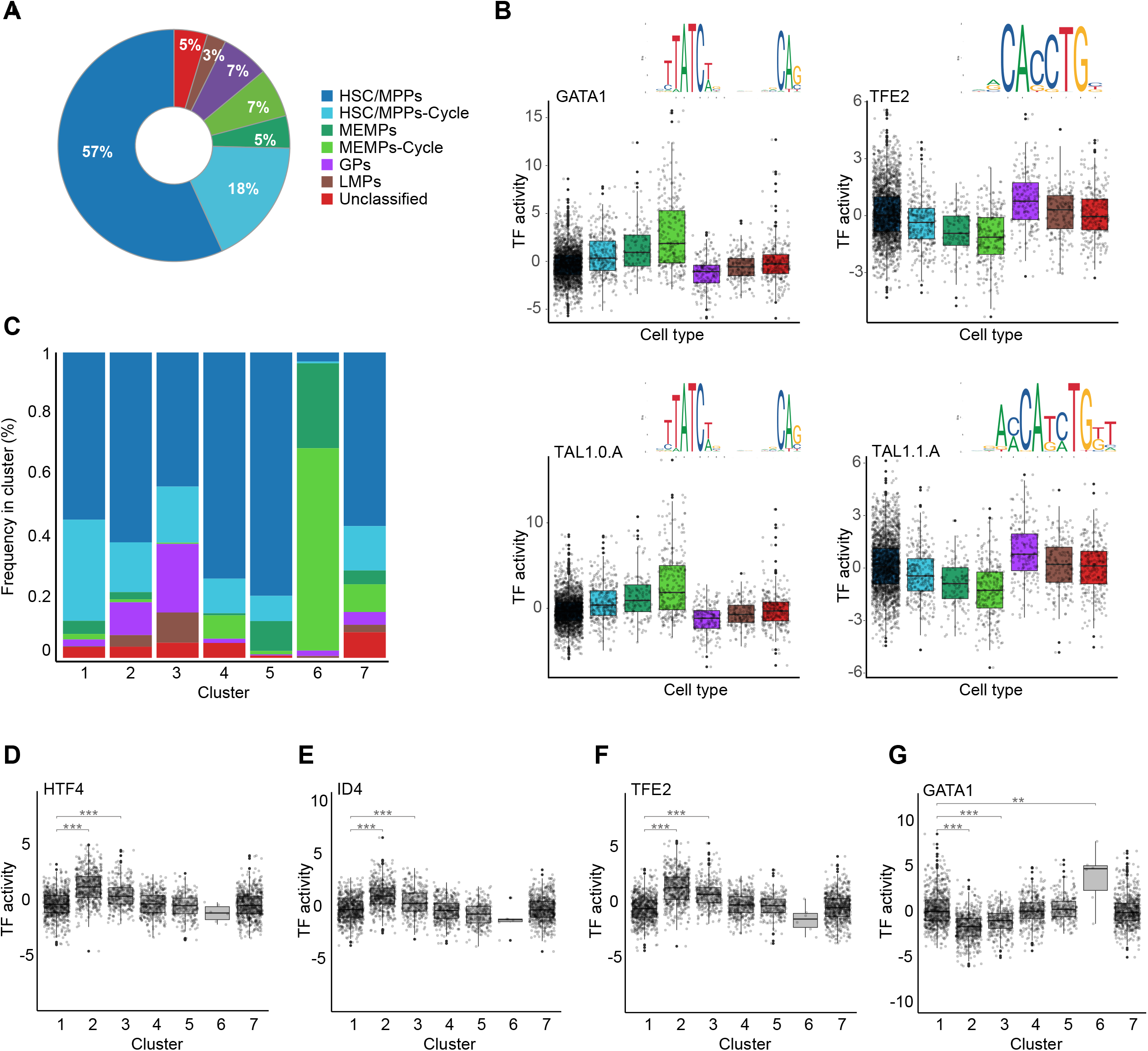
Integration of scRNA-Seq and scATAC-Seq data. **A.** Donut plot showing the percentage of scATAC-Seq cells automatically assigned to different cell types. **B.** Boxplot showing the accessibility of GATA1, TFE2, TAL1.0.A, and TAL1.1.A motifs in the annotated cell types. On the top right of each boxplot, the TF sequence logos from JASPAR database similar to the analysed motifs are shown. Cluster colours match those of **(A)**. **C.** Barplot showing the percentage of cells within each cluster assigned to the annotated cell types. Cluster colours match those in **(A)**. (D-G) Boxplots showing the accessibility of lineage-specific TF motifs in HSC/MPPs across the seven clusters. **D.** HTF4 (p-value_1,2_<2×10^-16^; p-value_1,3_<2×10^-16^). **E.** ID4 (p-value_1,2_<2×10^-16^; p-value_1,3_=10^-13^). **F.** TFE2 (p-value_1,2_<2×10^-16^; p-value_1,3_<2×10^-16^). **G.** GATA1 (p-value_1,2_<2×10^-16^; p-value_1,3_<2×10^-16^; p-value_1,6_=0.0071). p-value<0.001 (***); 0.001<p-value<0.01 (**); 0.01<p-value<0.05 (*); p-value≥0.05 (ns). Force-Directed Graph - FDG; ForceAtlas2 - FA2.

The frequency of assigned cell types in the scATAC-Seq data set was highly concordant with the ones from scRNA-Seq data (Supplementary figure 4B) suggesting that overall the two modalities, i.e., chromatin accessibility and transcriptome are correlated. To validate the cell type assignment of scATAC-Seq cells, we examined the accessibility of selected lineage-specific TF motifs in each of the annotated cell types (Figure 4B). In line with the predicted annotations, the GATA1 motif showed the highest accessibility in MEMPs and MEMPs-Cycle, whereas TEF2 (known to play a role in myeloid and lymphoid differentiation, (Miyamoto et al., 2002)) was most active in GPs and LMPs. Confirming our earlier observation, two distinct TAL1 motifs had anticorrelated accessibility. TAL1.0.A was preferentially active in MEMPs and MEMPs-Cycle, while TAL1.1.A in GPs and LMPs (Figure 4B).

The Force Atlas representation of the classified scATAC-Seq cells revealed, however, considerable intermixing of different cell types across the trajectory with enrichment of MEMPs/MEMPs-Cycle in cluster 6 and to a lesser extent of GPs and LMPs in clusters 2 and 3 (Figure 4C). HSC/MPPs(Cycle) were distributed across all seven clusters. This wide distribution of HSC/MPPs(Cycle) across multiple clusters within scATAC-Seq data suggested that, even though chromatin accessibility and the transcriptional state of foetal HSC/MPPs are correlated, there is extensive chromatin priming in the HSC/MPP population that results in their heterogeneity.

Next, we compared the accessibility of selected lineage-specific TF motifs in HSC/MPPs across the seven clusters (Figure 4D-G). We observed a low level of activity of all examined TFs in cluster 1 followed by a statistically significant increase of HTF4, ID4, and TFE2 and decrease of GATA1, in HSC/MPPs in clusters 2 and 3. GATA1 activity, however, increased in HSC/MPPs in cluster 6. Our data suggest that, within the transcriptionally homogeneous population of HSC/MPPs, there are significant differences in the activity of specific TFs that may precede gene expression and mark initial priming of HSC/MPPs prior to their commitment to the specific lineage. To explore further this “time lag” between the chromatin accessibility and gene expression during differentiation we examined scRNA-Seq and scATAC-Seq data for the top GATA1-regulon target genes (ranked based on the AUCell score) identified by pySCENIC (Figure 5). We looked at the accessibility of both gene promoters (+/− 3 kb from TSS) and distal regulatory regions (+/− 50 kb from the TSS) as well as the expression levels of the selected target genes along the MEMP differentiation trajectory (Figure 5A). We observed that promoters of GATA1-regulon target genes were often open in HSC/MPPs prior to any noticeable gene expression (Figure 5A). Thus, in line with our previous observation, chromatin accessibility in HSC/MPPs preceded transcriptional changes that were only present in more differentiated cells. Interestingly, promoter accessibility of GATA1 target genes was overall lower in cluster 6 (MEMPs) compared to cluster 1 (HSC/MPPs) (Figure 5B, D, E), and coincided with lower promoter co-accessibility of the antagonistic genes (i.e. genes that are specific for distinct lineages), (Figure 5F). In contrast, the accessibility of distal regulatory elements/enhancers had higher accessibility in cluster 6 compared to cluster 1 (Figure 5C). This may indicate that in GATA-regulon genes may be primed at promoters while it is the enhancers that contribute the cell-type-specific expression.

**Figure 5.**
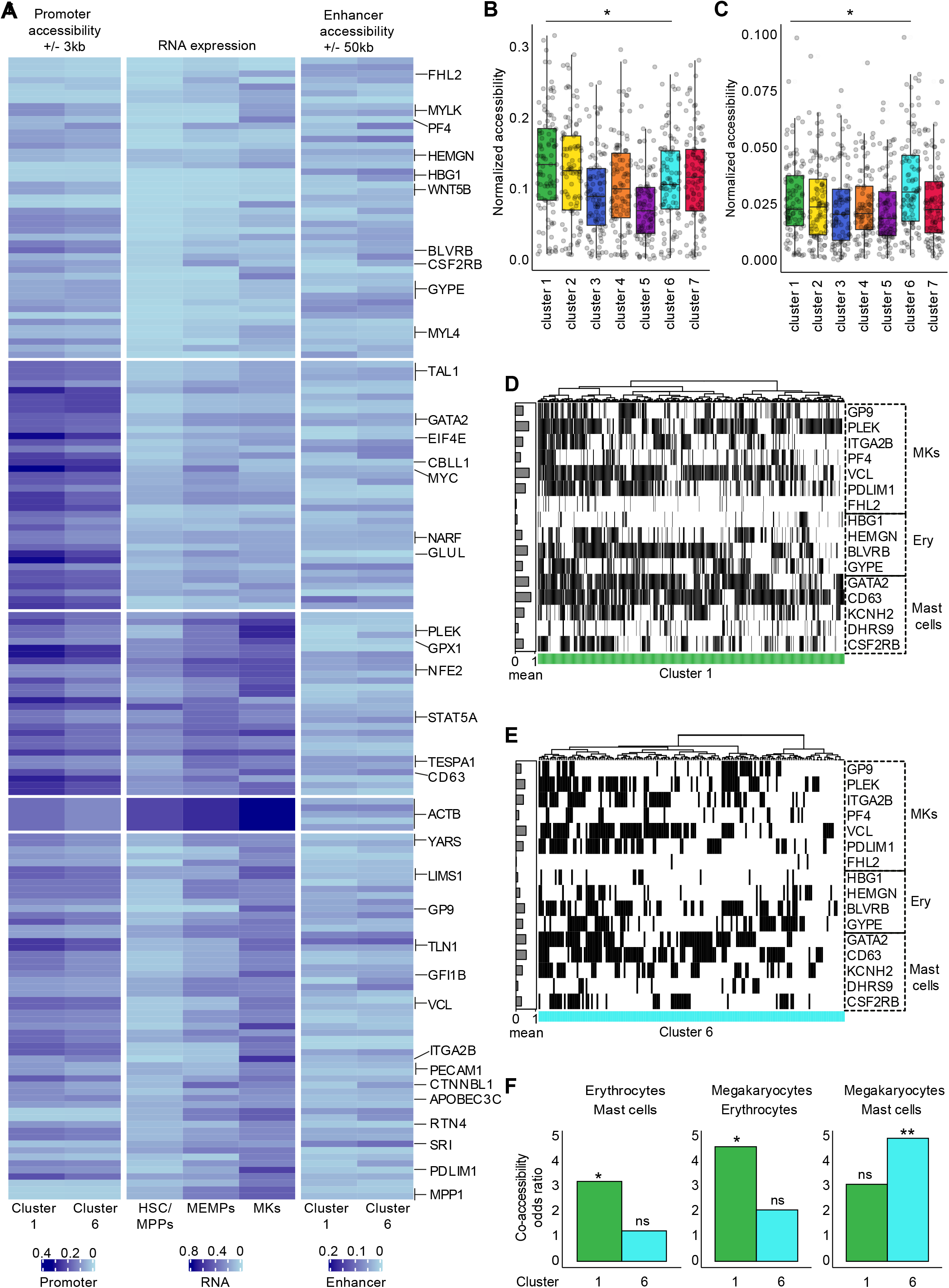
Chromatin accessibility and expression dynamics of GATA1-regulon target genes. **A.** Heatmap showing the chromatin accessibility changes for promoters (left) and related distal regulatory elements (right), as well as RNA expression (centre) for the target genes of GATA1 regulon obtained from pySCENIC. Only target genes with importance higher than 4 were considered. We normalised the expression of the set of GATA1 target genes into the range [0, 1]. For the promoters, the reads that overlapped the TSS regions +/−3 kb were extracted, normalised and scaled into the range [0, 1] for each cell. To identify the enhancers for each gene, we took peaks around TSS +/−50 kb (excluding +/−3 kb region), which have predicted GATA1 binding sites within them. We used normalized values for such peaks and scaled values from 0 to 1. Resulted values were summarized using mean per each cluster. For the visualisation purposes, we pooled all data together and clustered with 5 centroids using k-means. **B-C.** Boxplot showing the difference in chromatin accessibility for GATA1-regulon genes for all identified scATAC-Seq clusters. Values were obtained following similar criteria as described in **A**. **B.** Promoters. Significant p-values: p-value_1,3_=7.7×10^-5^, p-value_1,4_=0.005, p-value_1,5_=3×10^-9^, p-value_1,6_=0.026, p-value_2,3_=0.001, p-value_2,4_=0.047, p-value_2,5_=1.8×10^-7^, p-value_3,5_=0.023, p-value_3,6_=0.034, p-value_3,7_=0.018, p-value_4,5_=3.4×10^-4^, p-value_5,6_=8.7×10^-6^, p-value_5,7_=5.5×10^-6^.**C.** Enhancers. Significant p-values: p-value_1,6_=0.017, p-value_3,6_ =0.006, p-value_3,6_=2.9×10^-4^, p-value_4,6_=9.1×10^-4^, p-value_5,6_=1.3×10^-4^, p-value_6,7_=0.004. **D-E**. Heatmaps of the binarised chromatin accessibility in cluster 1 (D) and cluster 6 (E) for the promoters (+/−3 kb from TSS) of the selected marker genes of Megakaryocytes (MK), Erythrocytes (Ery), and Mast cells. Barplots on the left side of the heatmaps show the mean accessibility of the gene promoter for each cluster. **F.** Barplots showing the co-accessibility of promoters’ of lineage-specific marker genes. Fisher exact test was used to check if binarised accessibility of promoters from different marker genes is associated between each other. Odds ratios of Fisher’s exact tests are reported on the y-axes. P-value (Mast-Ery: cluster 1) = 0.03149, p-value(Mast-Ery: cluster 6) = 0.6379; p-value(MK-Ery: cluster 1) = 0.01026; p-value(MK-Ery: cluster 6) = 0.1734; p-value (MK-Mast: cluster 1) = 0.3007; p-value (MK-Mast: cluster 6) = 0.005428.

### Validation of HSC/MPP identity and their differentiation capacity

Given the observed limitation of commonly used sorting markers to isolate pure progenitor populations, we devised a new FACS sorting strategy for HSC/MPPs based on cell-surface markers selected from the top 20 marker genes for this cluster in our scRNA-Seq dataset. The refined panel for HSC/MPPs included Lin- CD34+ CD38− CD52+ CD62L+ CD133+ (CD-REF from now on, Figure 6A).

**Figure 6.**
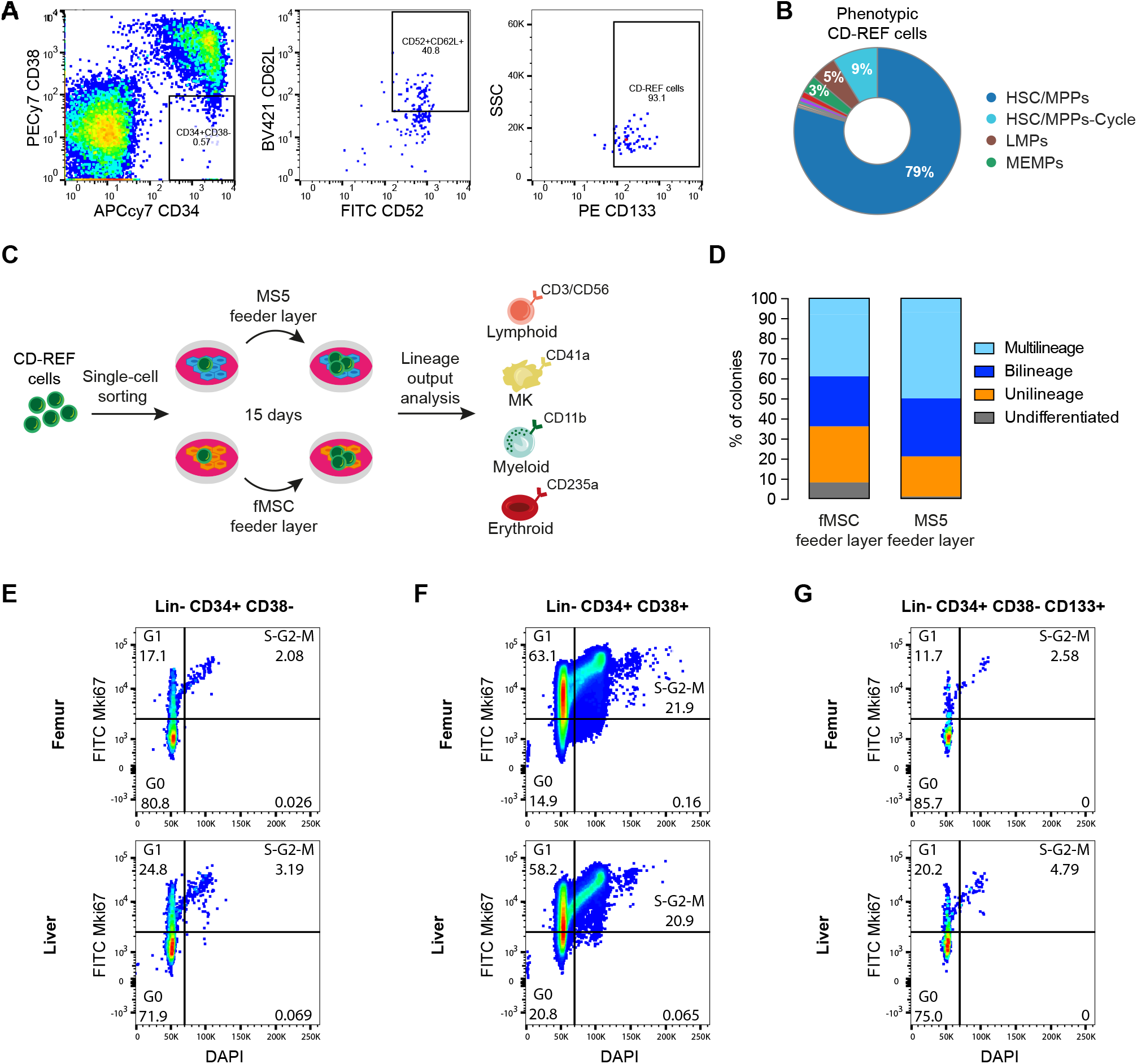
Refining the sorting strategy to isolate foetal HSC/MPPs. **A.** Novel FACS panel (CD-REF panel) designed to increase the purity of the sorted HSC/MPP population. After excluding debris, doublets, and Lin+ cells, CD34+ CD38− CD52+ CD62L+ CD133+ were sorted. **B.** Donut plots showing the percentage of transcriptionally defined (i.e., manually curated) cell populations in the phenotypically defined CD-REF population. The colours correspond to the identified cell types. **C.** Schematic overview of the single-cell *in vitro* differentiation assay. Single CD-REF cells were sorted in liquid culture with either a mouse stromal cell line (MS5) or a human foetal primary feeder layer (fMSC). After 15 days of culture, lineage output was assessed by the expression of lineage markers CD41a (megakaryocytic), CD235a (erythroid), CD3/CD56 (lymphoid), and CD11b (myeloid) by flow cytometry. **D.** Percentage of colonies derived from single CD-REF cells characterised by quadrilineage, trilineage, bilineage, unilineage and undifferentiated lineage output, on two different feeder layers (*n*=201 colonies, *n*=2 foetuses per feeder layer). **E-G.** Representative flow-cytometric images of cell-cycle analysis by Mki67/DAPI co-staining of CD34+CD38− (E), CD34+CD38+ (F) and CD34+CD38−CD133+ (G) in matched foetal liver and femur.

We FACS sorted cells from the femur BM using the CD-REF panel and profiled them again by scRNA-Seq and single-cell *in vitro* differentiation assays. CD-REF cells on average accounted for 40% (± 13%, *n*=4) of Lin- CD34+ CD38− cells in the femur, based on FACS analysis. The scRNA-Seq analysis of cells sorted with the refined panel showed that ~88% of CD-REF cells labelled HSC/MPP and HSC/MPP-Cycle clusters combined (Figure 6B) compared to commonly used CD panels for HSCs (Lin- CD34+ CD38− CD45RA- CD90+ CD49f+) and MPPs (Lin- CD34+ CD38- CD90- CD45RA- CD49f- CD10- CD7-) where ~59% and ~73% respectively of sorted cells had a transcriptional signature of our most immature cell population (Figure 1C, Supplementary figure 4B).

To assess the differentiation potential and robustness of the lineage output of CD-REF cells we sorted individual cells from three foetuses on either mouse MS5 feeder layer or on a more physiologically relevant, primary human foetal mesenchymal stem cells (fMSCs) (Figure 6C, see Methods section). After two weeks, 80% of cells sorted on MS5 and 85% of cells sorted on human foetal fMSCs generated colonies. In total, we analysed 201 colonies for their size and lineage output (erythroid-Ery, myeloid-My, megakaryocytic-Mk, lymphoid-Ly) using FACS (see Methods and Supplementary figure 7A). Our FACS analysis revealed that 7% of colonies on MS5 and 8% on fMSCs were quadri-lineage, 43% and 31% were tri-lineage, 29% and 25% were bilineage, 20% and 28% were unilineage and 1% and 8% were undifferentiated colonies, respectively (Figure 6D).

Next, we sorted individual CD-REFs and immunophenotypic HSCs (CD34+CD34-CD90+CD45RA-CD49f+/−) from the bone marrow and liver of the same foetus (*n*=2) on MS5 feeder layer and assessed 324 cells in total for their lineage output. Our analysis showed that the liver and femur derived CD-REF cells had comparable efficacy of colony formation and the lineage output, suggesting that CD-REF enriches for the population of cells with multilineage output in both foetal liver and bone marrow. Similarly, the lineage output of CD-REF cells and immunophenotypic HSC was comparable, however, the efficacy of colony formation appeared to be higher in CD-REF vs phenotypic HSCs isolated from the femur (Supplementary figure 7). Our finding that CD-REF cells indeed have multipotent potential and the lineage output comparable to phenotypic HSCs is in line with our observation that these cells sit at the tip of the differentiation trajectories. All together, we computationally and functionally confirmed that CD-REF represents a highly enriched population of HSC/MPPs.

### Comparative analysis of HSC/MPP cells from different haematopoietic organs

Cells in the HSC/MPP cluster originated from the liver, femur, and hip. This provided a unique opportunity to assess potential qualitative and quantitative differences in the HSC/MPP population that originated from foetal liver or bone marrow. We first applied Fisher's exact test on the number of liver and femur cells in the different cell cycle states to determine whether there are non-random associations between the cycle state and the organ of origin (see Methods for further details). Interestingly, there was a statistically significant difference (p-value=4.25×10^-9^) in the cell cycle state of cells in the HSC/MPP cluster between femur and liver (Figure 7A). Cells in the femur were predominantly in G1 (~70% of cells) compared to the same population in the liver (~52%) (Figure 7B). These data suggest that HSC/MPPs become more quiescent as they migrate from the liver to the bone marrow during the second trimester of human development. In line with this, HSC/MPP cells were significantly less frequent in femur compared to the liver (Figure 7D), as confirmed by Fisher’s exact test on the total number of liver and femur cells (Figure 7E). This is in agreement with the increased proportion of phenotypic non-committed progenitors (CD34+ CD38−) found in the liver compared to the bone marrow (Figure 7C). Using Mki67 and DAPI staining we quantified the proportion of cells in the different stages of the cell cycle: G0 (Mki67-DAPI-), G1 (Mki67+DAPI-), S-G2-M (Mki67+DAPI+), as previously described (Kim and Sederstrom, 2015). Our analysis showed that the CD34+CD38- population is less cycling in both foetal liver and femur compared to CD34+CD38+ population (Figure 6E-F). Furthermore, we further showed that the vast majority of CD-REF cells are in G0/G1 in both femur and liver but that nearly twice as many cells are in S-G2-M in the liver compared to the femur (Figure 6G).

**Figure 7.**
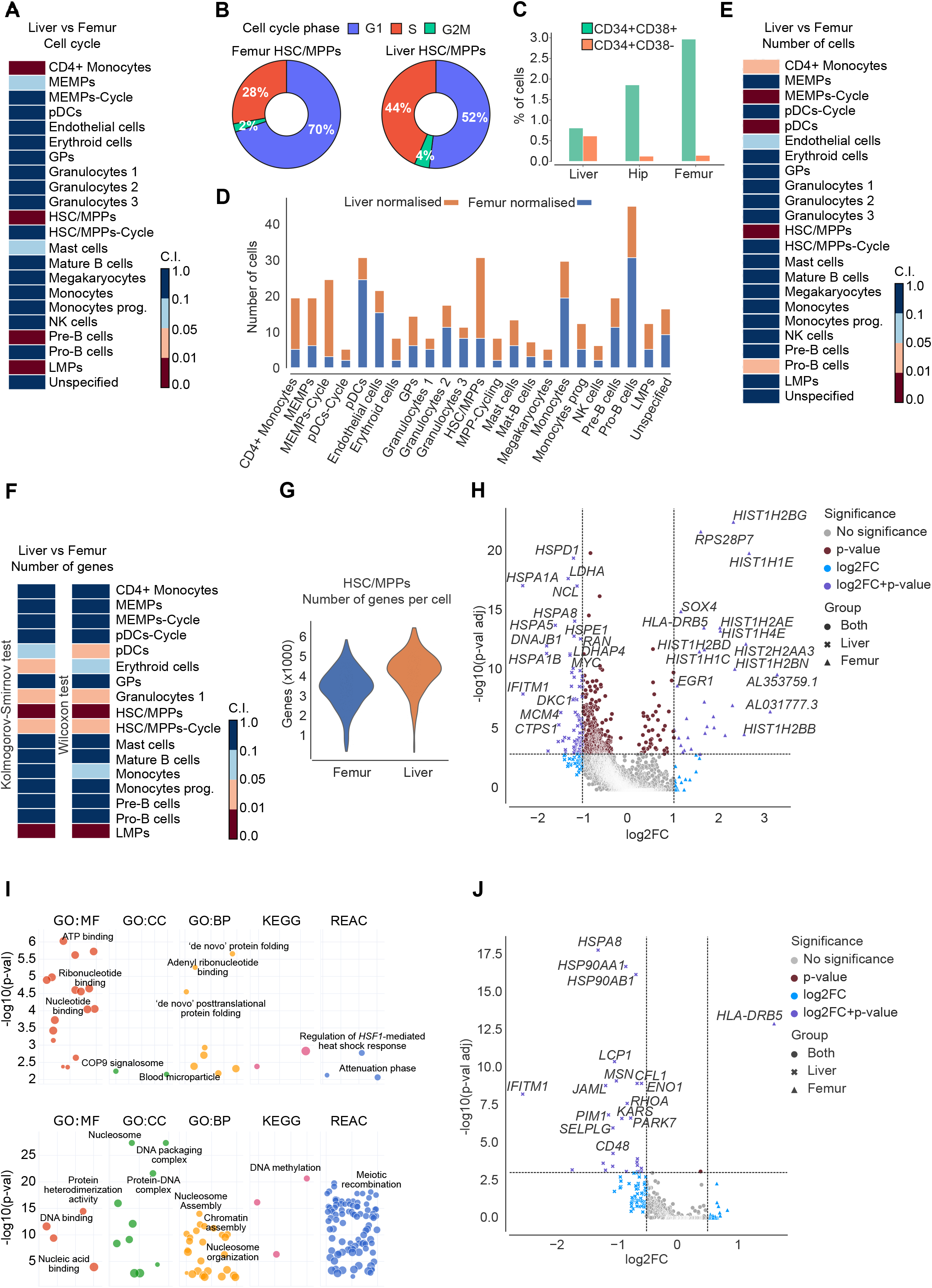
Statistically significant differences between femur and liver cells across cell types. **A.** Heatmap showing the confidence interval of Fisher's exact test on the normalised number of different haematopoietic cell types sorted from the liver and femur in G2M/S compared to G1. **B.** Donut plots displaying the percentage of cells in G1, G2M, and S in HSC/MPPs sorted from femur or liver. **C.** Bar plot representing the proportion of CD34+ CD38− and CD34+ CD38+ cells of total live cells present in the liver and bone marrow (femur and hip) *n*=15. **D.** Bar plot of the normalised distributions of the number of cells in each cell type sorted from liver or femur. **E.** Heatmap showing the confidence interval of Fisher's exact test on the normalised number of cells in each cell type collected from liver or femur **F.** Heatmaps depicting the confidence interval of the KS (left) and MWW (right) test on the number of expressed genes in each cell type collected from femur or liver cells. Notice that all the confidence intervals are split into 4 subintervals (i.e., [0, 0.01] strong statistically significant difference; (0.01, 0.05] statistically significant difference; (0.05, 0.1] marginal statistically significant difference; (0.1, 1] no statistically significant difference). **G.** Violin plot of the number of expressed genes in HSC/MPPs collected from the femur (blue) or liver (orange). **H.** Volcano plot showing DEGs in HSC/MPPs collected from femur or liver cells. The x-axis shows the *log2* fold-change (magnitude of change), while the y-axis shows the −log10 adjusted p-value (statistical significance). We used the Wilcoxon rank-sum with the Benjamini-Hochberg correction. Colours represent the significance of the genes, both in terms of p-value and *log2* fold-change. **I.** Bubble plots showing the top Gene Ontology terms (GO:MF - Molecular Function, GO:CC - Cellular Component, GO:BP - Biological Process), Kyoto Encyclopedia of Genes and Genomes, and Reactome calculated by using the DEGs in HSC/MPPs collected from the liver (top) vs femur (bottom) cells. **J.** Volcano plot showing DEGs in HSC/MPPs collected from femur or liver cells by considering only genes that encode plasma membrane proteins. The x-axis shows the *log2* fold-change (magnitude of change), while the y-axis shows the −log10 adjusted p-value (statistical significance). We used the Wilcoxon rank-sum with the Benjamini-Hochberg correction. Colours represent the significance of the genes, both in terms of p-value and *log2* fold-change.

In order to evaluate if there is a statistically significant difference in the number of expressed genes between HSC/MPPs collected from the liver and femur, we used both the Kolmogorov-Smirnov (KS) and Mann-Whitney-Wilcoxon (MWW) test. We applied a subsampling strategy to downsample the cluster with more cells and balance the two distributions (see Methods). KS and MWW revealed a statistically significant decrease in the number of expressed genes in HSC/MPPs in the femur compared to the liver (Figure 7F-G). Gene-set enrichment analysis, using pathway databases, of differentially expressed genes between the liver and femur revealed that HSC/MPPs in the femur up-regulate genes involved in nucleosome assembly, chromatin assembly, and DNA packaging such as *HIST1H1E* and *HIST1H2BN*, possibly marking their entry into quiescence (Figure 7H). Interestingly, DE analysis of genes that encode membrane proteins revealed statistically significant upregulation of genes related with actin cytoskeleton remodelling, cell adhesion, and migration (e.g., *JAML*, *SELPLG*, *LCP1*, *MSN*, *RHOA*) in HSC/MPPs in the liver compared to the femur (Figure 7I). This would be in line with the higher propensity of liver HSC/MPPs to migrate to other tissues such as bone marrow. In addition, we detected higher expression of interferon-induced gene *IFITM1* in foetal liver, known to play a role in the transduction of antiproliferative and adhesion signals (Figure 7J). This shift of HSC/MPPs from highly proliferative to quiescent as well as downregulation of genes involved in actin cytoskeleton remodelling, as they migrate from foetal liver to bone marrow, signifies the role of the niche in the modulation of HSC/MPPs behaviour.

## Discussion

Here, we present an integrative analysis of the single-cell transcriptome and chromatin accessibility of human foetal HSPCs. Our strategy involved plate-based sorting of well-defined immunophenotypic HSPCs from matched foetal liver and bone marrow. This approach enabled us to go beyond cataloguing heterogeneity of cellular states during foetal haematopoiesis and to: *i)* examine the extent to which phenotypic markers used over the last decade coincided with the true nature of the sorted foetal blood populations, *ii)* refine the sorting strategy for HSC/MPPs, *iii)* identify cell-cycle and gene expression differences between HSC/MPPs from foetal liver and bone marrow, *iv)* infer HSPCs differentiation trajectory, and *v)* explore lineage priming within the HCS/MPP population.

In doing so, we observed a striking level of heterogeneity in all immunophenotypic HSPCs, with more than ten transcriptionally-defined cell populations identified in each of the progenitor populations. Although this is consistent with previous studies of human adult and cord blood haematopoiesis (Knapp et al., 2018), it further emphasized the need for refining the sorting strategy for human foetal HSPCs. Our CD-REF panel achieved nearly 90% enrichment of HSC/MPPs, which we validated using single-cell *in vitro* differentiation assays and scRNASeq. CD-REF cells comprised 40% of all CD34+ CD38− cells in the foetal bone marrow with the majority of HSC/MPPs not cycling. The shift from highly proliferative state to quiescence coincided with the migration of HSC/MPPs from the foetal liver to bone marrow suggesting an important role of the niche in the modulation of HSC/MPPs behaviour. This is remarkably different from previous studies in mice, where an extensive proliferation of HSPCs in the bone marrow continued up to three weeks after birth (Bowie et al., 2006).

Downstream of the HSC/MPPs, we identified three highly proliferative oligopotent progenitor populations (MEMPs, LMPs, and GPs). Integrative scRNA-Seq and scATAC-Seq analysis of HSC/MPPs and all main progenitor populations revealed a correlation between chromatin accessibility and gene expression but also pointed out that, within transcriptionally homogeneous HSC/MPPs, there are multiple subpopulations that differed in their overall chromatin accessibility as well as lineage-specific TF activity. This indicates that within the HSC/MPP population, regulatory programmes permissive for different fates are being primed on the chromatin level, prior to their commitment to a specific lineage. The higher coordination of transcription and chromatin accessibility only occurred along the commitment of HSC/MPPs towards MEMPs, implying a hierarchy of different levels of commitment in the foetal progenitor compartment with MEMP population being the most committed (compared to LMPs and GPs). Overall, our study has provided a high resolution transcriptional and chromatin accessibility map of foetal HSPCs from the liver and bone marrow that will be essential for further exploration of HSC/MPPs in the context of blood pathologies and for the purpose of regenerative medicine.

### Limitations of the study

In this study, we characterised human foetal liver and bone marrow haematopoiesis using a combination of single-cell transcriptomics/epigenetics and in vitro single-cell differentiation assays. In order to avoid perturbations caused by freezing and thawing cycles, all experiments were performed on freshly-isolated tissues. Such experimental design and the nature of analysed tissues come with a few limitations: 1) samples are rare and 2) the cellularity varies significantly between different stages of development and individual foetuses, especially in the bone marrow. As a result, the number of cells available for the analysis was limited. For this reason, we were not able to obtain enough cells to perform xenotransplantation experiments to confirm the stem cell identity and self-renewing potential of our CD-REF cells, collected from bone marrow. Instead, we used single-cell in vitro assays as an alternative, albeit not optimal, readout of the multilineage potential of a cell. In addition, we could only collect a limited number of distinct phenotypically defined populations from individual foetuses and, therefore, for any given population, the number of analysed samples was relatively low.

## Acknowledgements

The authors would like to thank the WTSI Cytometry Core Facility for their help with single-cell index sorting and the WTSI DNA pipelines and the CRUK Cambridge Institute Genomics Core Facility for their contribution in sequencing the data. We would also like to thank the CellGen bioinformatics team for their help with pre-processing of scRNA-Seq data and Jana Eliasova for her precious support and help with the illustrations. We would like to thank Alice Martin for her help with pySCENIC analysis. Finally, we would like to thank the Human Developmental Biology Resource (HDBR) for providing samples.

The study was supported by European Research Council project 677501 - ZF_Blood (to A.C. and A.M.R), EMBO small grant (to A.C.) and a core support grant from the Wellcome Trust and MRC to the Wellcome Trust - Medical Research Council Cambridge Stem Cell Institute.

## Authors contributions

A.C. and A.M.R. conceived the study; A.M.R. performed all the experiments with help from B.M., P.M.S., J.X., and E.P.; A.T. carried out the computational analysis of the scRNA-Seq data; S.G.R. carried out parts of the computational analysis, under the supervision of A.T.; I.B. performed the analysis of the scATAC-Seq data and the integrative analysis of scATAC-Seq and scRNA-Seq, under the supervision of J.B.Z.; I.M. carried out the statistical analysis of *in-vitro* experiments; A.C., A.M.R, and A.T. designed the figures and wrote the manuscript with inputs from the other authors. All authors approved the final version of the manuscript.

## STAR Methods

### RESOURCES AVAILABILITY

#### Lead Contact

Further information and requests for resources and reagents should be directed to and will be fulfilled by the Lead Contact, Dr Ana Cvejic.

### Materials Availability

This study did not generate new unique reagents.

### Data and Code Availability

The raw RNA-Seq data (i.e., fastq files) and cell assignment are deposited at ArrayExpress with accession code E-MTAB-9067, while the raw ATAC-Seq data (i.e., fastq files) and cell assignment are deposited at ArrayExpress with accession code E-MTAB-9068.

All scripts, functions, and Jupyter Notebook developed for this study are freely available on GitLab: https://gitlab.com/cvejic-group/integrative-scrna-scatac-human-foetal. The repository also contains the gene expression and fragment matrices.

## EXPERIMENTAL MODEL AND SUBJECT DETAILS

### Ethics and Tissue acquisition

Human foetal bone and liver samples were obtained from 33 foetuses aged 17-22 pcw, following termination of pregnancy and informed written consent. The human foetal material was provided by the Joint MRC/Wellcome Trust (Grant MR/R006237/1) Human Developmental Biology Resource (http://www.hdbr.org), in accordance with ethical approval by the NHS Research Health Authority, REC Ref: 18/LO/0822.

## METHOD DETAILS

### Tissue processing

Tissues were kept in cold DMEM medium (Invitrogen) until dissection and processed on the same day of collection. Single-cell suspensions were generated from matched foetal liver and bone tissues after rinsing them with cold PBS (Gibco). Liver samples were passed through a 70 μm strain into a falcon tube prefilled with cold PBS. Bone marrow from long bones was isolated by flushing cold PBS into the diaphysis and collected into a falcon tube. Bone marrow from hip bone was collected by dissecting the bone with a sterile scalpel and flushing cold PBS in the marrow cavity into a falcon tube. The suspension obtained from long bones and hip bones was then passed through a 70 μm strain into a new falcon tube. Cells were then centrifuged for 5 minutes at 300 g, 4°C and the pellet was resuspended into the RBC lysis buffer (eBioscience) for 2 minutes at room temperature, after which 20 ml of cold PBS were added to stop the lysis reaction. RBC step was not performed when sorting erythroid cells. Live cell enrichment was performed using MACS columns (Miltenyi Biotec - 130-090-101) following the manufacturer's instructions. When sorting CD34+ or CD45+ cells, column enrichment was performed using MACS columns (Miltenyi Biotec - 130-046-702 and 130-045-801 respectively for CD34+ and CD45+ cells), following the manufacturer's instructions.

### Fluorescence-activated cell sorting

Cells were stained with antibody cocktails in a total volume of 100 μl 5% FBS (Gibco) in PBS for 30 minutes at 4°C, centrifuged for 5 minutes at 300 g, 4°C, resuspended in a final volume of 500 μl of 5% FBS in PBS and subsequently filtered into polypropylene FACS tubes (ThermoFisher). For scRNA-Seq experiments, single cells were index sorted using a BD Influx Sorter into wells of 96-well plates (4titude) prefilled with 2 μl of lysis buffer consisting of 0.2% Triton X-100 (Sigma) and 1 U/μl RNAse inhibitor (Life Technologies) in nuclease-free water (Invitrogen). For scATAC-Seq experiments, 5,000 - 20,000 cells were sorted using a BD Influx machine into 1.5 ml tubes (Eppendorf). Following bulk tagmentation with Tn5 (Chen et al., 2018), single nuclei were index sorted in wells of 384-well plates (Eppendorf) prefilled with 2 μl of lysis buffer consisting of 0.2% SDS, 20 μg/ml proteinase K (Ambion), 50 mM Tris-HCl (Gibco) and 50mM NaCl (Sigma) in nuclease-free water.

### Library preparation

The Smart-Seq2 method (Picelli et al., 2014) was used for library preparation for the scRNA-Seq experiments, with some modifications as described in (Macaulay et al., 2016). The quality of libraries was evaluated with Bioanalyzer (Agilent). Good-quality libraries were subsequently quantified with KAPA Library Quantification Kit (Roche) and submitted for sequencing. Library preparation for the scATAC-Seq experiments was performed using a recently described method (Chen et al., 2018). Library traces were evaluated using Bioanalyzer.

### Sequencing

Libraries for scRNA-Seq experiments were multiplexed using Nextera Index sets A, B, C, and D (v.2, Illumina) and sequenced on HiSeq4000 and NovaSeq6000 (Illumina) in pair-end mode, with an interquartile range (IQR) of 697,427 uniquely mapped reads (average: 666,632; standard deviation: 557,274). Libraries for scATAC-Seq experiments were sequenced on HiSeq4000 in pair-end mode, with a mean read count of 473,886 and IQR 341,210.

### Upstream analysis of scRNA-Seq data

Smart-Seq2 sample demultiplex fastq files were quality checked, aligned and quantified by using the scRNA-Seq pipeline. This pipeline is based on STAR with default parameters (v.2.5.4a) (Dobin et al., 2013) index and annotation from the Ensembl release 91 of the GRCh38 human reference genome. Transcript and gene counts were quantified using the option *quantMode GeneCounts* provided by STAR. Since we used different sets of well-defined antibodies to isolate different cell types, we applied specific thresholds for each sample to filter out both the cells and genes (Supplementary table 3). We detected on average 3,642 genes per cell (IQR: 2,239; standard deviation: 1,621).

### Downstream analysis of scRNA-Seq data

In what follows, for each function that we applied, we only report the parameter settings we modified. All other parameter settings of the functions are the default ones provided by the used computational libraries. We performed the downstream analysis of scRNA-Seq using the Python (v.3.6.9) package SCANPY (v.1.4.5.1) (Wolf et al., 2018). Our pipeline included: 1) a QC step (number of identified counts and number of expressed genes using the *filter_cells* function, and the fraction of mitochondrial genes). We obtained the 4,504 cells that were used in the next steps (Supplementary table 2), 2) removing the genes expressed in less than 10 cells (*filter_genes* function), 3) data normalisation (*normalize_per_cell* function with scaling factor 10,000 and *log1p* function), 4) detection of the top 1,000 highly variable genes (HVGs) (*highly_variable_genes* function, in which the HVGs were selected separately within each batch and then merged, where each batch corresponds to a specific sample), 5) scaling of the features to unit variance and zero mean (*scale* function with max_value equal to 10), 6) application of scAEspy on the HVG space by considering the raw expression (i.e., counts) (Tangherloni et al., 2019), 7) batch correction by sample applying BBKNN algorithm (v.1.3.6, *bbkkn* function with use_faiss equal to false, approx equal to false and the Euclidean distance) to the latent space (16 components) generated by the used AE, 8) Leiden algorithm (*leiden* function with resolution equal to 2.2) applied to the neighbourhood graph generated by BBKNN. The 27 obtained clusters were manually annotated by considering the merged data using well-known cell-type specific genes and the Differentially Expressed Genes (DEGs). DEGs were computed by using *rank_genes_groups* function (Wilcoxon rank-sum with adjusted p-values for multiple testing with the Bonferroni correction), which compares each cluster to the union of the rest of the clusters. The clusters that either did not express specific cell type genes or expressed marker genes of different cell types had been iteratively subclustered. Specifically, we applied the Leiden algorithm (*leiden* function with resolution equal to 0.5) to subcluster Endothelial cells, obtaining four distinct clusters: the first two clusters have been annotated as Monocytes 2, the third as NK cells and the fourth as Endothelial cells. Finally, we used the Leiden algorithm (*leiden* function with resolution equal to 0.5) to cluster the Unspecified cluster getting four clusters. We merged three clusters with the HSC/MPP cluster while one was annotated as Unspecified.

### Dimensionality reduction of scRNA-Seq data

After the detection of the first 1,000 HVGs, we applied scAEspy to HVG space by setting alpha and lambda equal to 0 and 2, respectively, in order to obtain the Gaussian Mixture Maximum Mean Discrepancy Variational AE (GMMMDVAE) (Tangherloni et al., 2019). We run GMMMDVAE for 100 epochs with a batch size equal to 100, one hidden layer of 64 neurons, a latent space of 16 neurons, 15 Gaussian distributions, learnable prior distribution, constrained Poisson loss function, and sigmoid activation function. Then, we applied BBKNN to the latent space (16 components) to generate the neighbourhood graph by identifying top neighbours of each cell in each batch separately. We applied UMAP (v.0.3.10, SCANPY *umap* function with random_state equal to 8 and n_components equal to 3) to the obtained neighbourhood graph.

### Trajectory analysis of scRNA-Seq data

In order to perform a detailed comparison among different trajectory modelling tools, the Dynverse tool (Saelens et al., 2019) was used. Based on the scoring system provided by Dynverse and following a careful inspection of the generated trajectories, we applied PAGA and Force-Directed Graph (FDG) to infer the development trajectories. We removed the endothelial cells and recalculated the neighbourhood graph (*neighbors* function with n_neighbors equal to 30) on the latent space (16 components) to exploit the data before batch correction (Luecken and Theis, 2019). We computed the PAGA graph (*paga* function with model equal to v1.2) and the ForceAtlas2 (FA2) using PAGA-initialization (*draw_graph* function, which exploits the FA2 class from fa2 (v.0.3.5) Python package, using the HSC/MPP cluster as root and maxiter equal to 1,000).

### Differential expression analysis

Following cluster annotation, we performed biologically-relevant pairwise DE tests between pairs of clusters to identify DEGs and to examine the quantitative changes in the expression levels between the clusters. Specifically, we tested MPPs against MPPs-Cycle, MEMPs against MEMPs-Cycle, MPPs against MEMPs, MPPs against LMPs, MPPs against GPs, and MEMPs against LMPs. In order to cope with the unbalanced distributions between two groups of cells, due to the different number of cells in each cluster, we used the following subsampling strategy. Given two groups of cells, the biggest group was randomly subsampled taking a number of cells equal to the number of cells composing the smallest group. For each gene, a two-sided T-test for the means of two independent samples (i.e., biggest group and subsampled one) was applied. We used the *ttest_ind* function (equal_var equal to false) provided by the Python SciPy (Virtanen et al., 2020) package (v.1.4.1). Since we did not assume that the two groups have identical variances, the Welch’s t-test was automatically applied. Then, we calculated the median of the p-values of these T-tests. We applied this subsampling strategy 1001 times and calculated the median of the medians to select the subset of the biggest group to run the DE analysis.

For a given subset of cells from the biggest group and the smallest one, we calculated the DEGs by applying the *rank_genes_groups* function (Wilcoxon rank-sum with adjusted p-values for multiple testing with the Benjamini-Hochberg correction). Then, we filtered out the obtained DEGs by using the *filter_rank_genes_groups* function (min_in_group_fraction equal to 0.3 and max_out_group_fraction equal to 1, so that a gene is expressed in at least 30% of the cells in one of the two tested groups; min_fold_change equal to 0). Following the aforementioned workflow, we compared cells from the liver and femur from the same cluster. Finally, we analysed HSC/MPPs and HSC/MPPs-Cycle to see which genes contributed to the observed difference between cells from femur and liver.

We also carried out a DE test to compare the expression of cell surface proteins in HSC/MPPs from femur and liver cells. As a first step, we selected 1) genes that encode CD molecules, 2) transmembrane genes available in CellPhoneDB (Efremova et al., 2020), and 3) genes that encode plasma membrane proteins from Uniprot (key KW-1003). Then, we applied the subsampling strategy comparing HSC/MPPs from femur and liver cells. Finally, we calculated the DEGs by considering only the genes that are expressed in at least 30% of the cells in one of the two tested groups.

### Differentiation pathway analysis

We performed a gene-set enrichment analysis, using pathway databases, comparing liver and femur HSC/MPP cells. Firstly, we calculated DEGs by comparing liver and femur applying the strategy described above. Then, we used g:Profiler (Raudvere et al., 2019) focusing on Gene Ontology (GO) terms, Kyoto Encyclopedia of Genes and Genomes (KEGG), and Reactome. Specifically, we applied the profile function provided by the Python *GProfiler* package for both liver and femur cells. As a query set, we used the liver (or femur) DEGs while as background we used the genes that are expressed in at least 30% of liver (or femur) cells. We also set the following parameters required by the *profile* function: *organism* equal to homo sapiens; *sources* equal GO terms, KEGG, and Reactome; *domain_scope* equal to custom_annotated; *significance_threshold_method* (i.e., the correction method for the p-values) equal to bonferroni; *user_threshold* (i.e., the threshold for the corrected p-values) equal to 0.01.

### Cell type classification

We trained both a Random Forest classifier (Pedregosa et al., 2011) and a DNN to predict the cell types by considering the top 5, 10, 20, 30, 50, and 100 marker genes for each cluster using the *log*-normalised expression. Since some marker genes are shared among the clusters, we considered them only once to avoid duplicated columns in the feature matrices. We merged the following clusters: HSC/MPPs and HSC/MPPs-Cycle as HSC/MPPs, MEMPs and MEMPs-Cycle as MEMPs, Granulocytes 1, Granulocytes 2, and Granulocytes 3 as Granulocytes; pDCs and pDCs-Cycle as pDCs; CD4+ Monocytes, Monocytes, and Monocyte Prog as Monocytes; Pre-B cells, Pro-B cells, and Mature B cells as B cells. Thus, we obtained 14 distinct clusters.

We used the *RandomForestClassifier* (n_estimators equal to 100 and Gini criterion) provided by Scikit-learn (Pedregosa et al., 2011) (v.0.21.2). We developed the DNN by using Keras^1^ (v.2.2.4) with Tensorflow (Abadi et al., 2016) (v.1.12.0) as backend. The network is composed of 2 dense hidden layers of 64 and 32 neurons, respectively. We added a dropout (50%) layer before the first layer as well as a dropout (30%) layer before the second layer. We trained the DNN for 1,000 epochs using the Adam optimiser (Kingma and Ba, 2014) by minimising the categorical cross-entropy loss function. We also set an early stopping with 100 epochs as patience to avoid overfitting.

We applied a stratified 10 fold cross-validation (Scikit-learn *StratifiedKFold* function) resampling procedure to evaluate both the Random Forest and DNN. The Random Forest achieved the best result when the top 30 marker genes per cluster were used (mean accuracy equal to 88.54% and standard deviation equal to 1.03%), while the DNN considering the top 30 (mean accuracy equal to 90.40% standard deviation equal to 1.31%) and 50 marker genes per cluster (90.25% and standard deviation equal to 0.98%).

As a further test, we evaluated the ability of our DNN to generalise on unseen data. We split the dataset into a train set (80%) and a test set (20%) (Scikit-learn *train_test_split* function with test_size equal to 0.2). We then divided the train set into a train set (85%) and a validation set (15%, (*train_test_split* function with test_size equal to 0.15). We trained our DNN with the train set, validating it using the validation set. When we took into account the top 30 marker genes, we achieved an accuracy equal to 91.50% on the validation set. When considering the top 50 marker genes the accuracy was 91.68%. Finally, we predicted the labels of the test set by obtaining an accuracy equal to 90.46% (30 marker genes) and 90.23% (50 marker genes).

### Upstream analysis of scATAC-Seq data

We performed the upstream analysis using the samtools (Li et al., 2009) (v1.9), bedtools (Quinlan, 2014) (v2.27.1), Picard tools^2^ (v2.9.0) and BWA (Li and Durbin, 2009) (v0.7.17). First, we aligned fastq files to the GRCh38 reference genome (average 473,886 reads per cell), followed by marking duplicates with *MarkDuplicates* function from Picard tools and removing duplicates using samtools *view* with -F 1804 parameter per each cell. Overall with average duplicates rate 77% we obtained 91,554 reads per cell after removing duplicates. Next, we transformed bam files to bed files using *bamtobed* bedtools function in bedpe mode and kept only fragments that are no bigger than 1000 bp using a custom script. We called peaks (for the clusters with more than 50 cells) using the SnapATAC approach (Fang et al., 2019) with macs2 (Zhang et al., 2008) parameters “--nomodel --shift 100 --ext 200 --qval 5e-2-B” and obtained 152,283 peaks. Importantly, for the downstream analysis in R, we binarized counts per cell using Signac^3^ *BinarizeCounts* function, resulting in 32,217 fragments per cell on average.

### Downstream analysis of scATAC-Seq data

The downstream analysis was done in R 3.6.1 applying Seurat (Butler et al., 2018) (Stuart et al., 2019) (v3.1.4), Signac (v0.2.4), chromVAR (Schep et al., 2017) (v.1.8) and Harmony (Korsunsky et al., 2019) (v1.0). The pipeline included a QC step (duplicates removal, number of fragments, fragments per peak, fraction of reads mapping to blacklist regions, nucleosome signal, and transcriptional start site (TSS) enrichment), application of LSI dimensionality reduction to the three samples independently (*RunTFIDF* function with method equal to 2, *FindTopFeatures* function setting min.cutoff to q0, and *RunSVD* function using the peaks as assay), batch correction by sample, lane, and organ applying Harmony on the first 50 LSI components, excluding the first one, (*RunHarmony* function setting assay.use to peaks, max.iter.harmony to 20, max.iter.cluster to 200, sigma to 0.25, and theta to 2, 4, 4 in order to weight more the batch related to samples). TF activities on the ATAC-seq data were calculated using the Signac implementation of chromVAR using the *RunChromVAR* function taking as tested motifs dataset from HOCOMOCO (Kulakovskiy et al., 2018) v11 human TF binding models database (769 TFs).

### Dimensionality reduction of scATAC-Seq data

We applied the UMAP algorithm to the first 50 LSI components corrected by Harmony (*RunUMAP* function with umap.method equal to uwot and n.neighbors equal to 10, *FindNeighbors* function setting annoy.metric to cosine). We identified seven distinct clusters by using the Seurat function *FindClusters* (resolution equal to 0.5).

### Trajectory analysis of scATAC-Seq data

We inferred the development trajectories by applying PAGA and FDG. We recalculated the neighbourhood graph using the SCANPY *neighbors* functions (n_neighbors equal to 30) on the 50 LSI components corrected by Harmony. We computed the PAGA graph (SCANPY *paga* function with model equal to v1.0) and used it to initialise the FA2 algorithm (SCANPY *draw_graph* function using cluster 1 as root and maxiter equal to 1,000).

### Integration of scRNA-Seq and scATAC-Seq data

We integrated scRNA-Seq and scATAC-Seq data using a recently developed method by Stuart *et al.* (Stuart et al., 2019). Namely, we used our scRNA-Seq data as reference dataset to train the classifier and automatically assign a cell type to each scATAC-Seq cell. The training of the classifier was performed using 511 CD34+ CD38- cells from our scRNA-Seq experiment. In order to have a suitable number of cells for each cell type to train the classifier, we considered scRNA-Seq clusters with at least 20 cells (i.e., HSC/MPPs, HSC/MPPs-Cycle, MEMPs, MEMPs-Cycle, GPs, and LMPs). We generated a gene expression matrix from our scATAC-Seq data set by assigning each peak to the gene by considering the genome coordinates of the gene body ± 3 kb. We applied the Seurat function *FindTransferAnchors* (query.assay equal to RNA_promoter, features equal to the counts of the RNA_promoter, and k.anchor equal to 6) on the Canonical Correlation Analysis (CCA) space because it was more suitable, compared to the LSI space, for capturing the shared feature correlation structure between scRNA-Seq and scATAC-Seq data. We assigned the cell types to the scATAC-Seq cells by applying the Seurat *TransferData* on the first 50 LSI components corrected by Harmony considering the calculated anchors (refdata equal to the six scRNA-Seq clusters). In order to avoid assignments based on a low score, all cells with the prediction score lower than 40% (the value of a uniform distribution of six clusters is 16,67%) were labelled as unknown.

### Transcription factor regulons prediction

To run SCENIC workflow on our raw scRNA-Seq data, we used an in-house constructed Snakemake pipeline via combining Arboreto package GRNBoost2 and SCENIC algorithms with default parameters. To predict transcription factor regulons, we used human v9 motif collection, as well as both *hg38 refseq-r80 10kb_up_and_down_tss.mc9nr.feather* and *hg38 refseq-r80 500bp_up_and_100bp_down_tss.mc9nr.feather* databases from the cisTarget (https://resources.aertslab.org/cistarget/). The resulting AUC scores per each cell and adjacency matrix were used for downstream analysis and visualization.

### Isolation of human foetal MSCs

Human primary fMSCs were isolated from the femur of a 19 pcw sample following an established protocol used for mouse bones (Perpétuo et al., 2019). Briefly, the bone was rinsed in PBS and the bone epiphyses cut with a scalpel. The bone marrow was flushed with 50 ml PBS, centrifuged at 300 g for 5 minutes, resuspended in alphaMEM medium (Thermo Fisher Scientific) supplemented with 2 mM L-glutamine (Thermo Fisher Scientific), 100 U/ml penicillin/streptomycin (Thermo Fisher Scientific) and 10% fetal bovine serum (Sigma) at a concentration of 5×10^6^ cells/ml and cultured at 37°C at 5% CO2. After 24 hours, floating cells were removed by washing twice with PBS and medium was changed twice a week until the culture was 70% confluent. Cells were cryopreserved until use.

### Single-cell in vitro culture

Single Lin- CD34+ CD38- CD62L+ CD52+ CD133+ cells, isolated from the foetal bone marrow of three different foetuses (20-22 pcw), were index-sorted into 96-well plates seeded with fMSCs or MS5 (obtained by DSMZ) and supplemented with cytokines as previously described (Velten et al., 2017). Cells were cultured for 15 days at 37°C at 5% CO2. At the end of the culture, colonies were filtered to exclude feeder layer cells, and their lineage output was assessed by the expression of CD41a (megakaryocytic-Mk), CD235a (erythroid-Ery), CD3/CD56 (lymphoid-Ly), and CD11b (myeloid-My) by flow cytometry using a BD LSR-Fortessa analyser. Colonies were considered positive for a lineage if ≥ 30 cells were detected in the relative gate.

### Cell cycle analysis

Cells from the foetal liver and the bone marrow were stained with cell-surface antibodies, fixed and permeabilised for 20 minutes at 4°C using the Cytofix/cytoperm kit (BD Biosciences). Cells were then stained with FITC-MKi67 antibody (BD Biosciences) overnight at 4°C and finally with DAPI prior to flow cytometry acquisition. Cell cycle phases were defined as follows: G0 (Mki67-DAPI-), G1 (Mki67+DAPI-), S-G2-M (Mki67+DAPI+).

## QUANTIFICATION AND STATISTICAL ANALYSIS

### Differences across cell types

In order to assess qualitative and quantitative differences between the haematopoietic cells collected from the liver and femur, we implemented different statistical tests. For each cluster, we calculated if there is a statistically significant difference in the number of cells (Test 1), the number of expressed genes per cell (Test 2), and the cell cycle state of blood cells collected from liver and femur (Test 3).

### Test 1

Since we used different gates to sort cells and we sorted a different number of cells in each experiment, we first normalised the number of cells from liver and femur. We selected only the matched gates (i.e., the gates where we sorted haematopoietic cells from both liver and femur). Then, we selected cells from the liver (or femur) from each gate in each of the clusters. For each cluster, we normalised the number of cells inside the cluster in the range [0, 100] by dividing the number of cells for the total number of cells of the gate in order to obtain a number of cells equal to 100. Next, for each cluster, we calculated the median of the cells in the liver (or femur) among the different gates. In order to evaluate if there is a statistically significant difference between the number of cells in the liver and femur considering all the clusters, we applied the ChiSq test by normalising the distributions (i.e, the median of the gates of each cluster) of the cells from liver and femur among the clusters. We applied the *chi2_contingency* function provided by the Python SciPy. Since the obtained p-value is equal to 1.02×10^-4^, we applied Fisher's exact test (SciPy *fisher_exact* function) to each cluster to find which clusters contributed to the difference.

### Test 2

In this test, we evaluated the number of expressed genes between cells collected from femur and liver. In order to remove possible technical effects for each cell, we divided the number of expressed genes by the number of reads uniquely mapped against the reference genome. For each cluster, we applied both the KS test (SciPy *ks_2samp* function) and the MWW test (SciPy *mannwhitneyu* function). Since the number of cells from femur and liver is very different in any given cluster (giving rise to unbalanced distributions) we used a subsampling strategy similar to that used for the DE analysis. We randomly subsampled the biggest group 1,001 times taking a number of cells equal to the number of cells composing the smallest group. We applied the KS (and MWW) test comparing the smallest group to the subsampled ones obtaining a distribution of p-values. Finally, we calculated the median of this distribution of p-values to evaluate if there is a statistically significant difference between the number of expressed genes in the cells from the liver and femur. Note that we excluded the clusters where the number of the cells from femur or liver was lower than 20.

### Test 3

For each cluster, we compared G2M/S and G1 states by normalising the number of cells from the liver and femur in the two states. We applied Fisher's exact test (SciPy *fisher_exact* function) to each cluster to find a possible statistically significant difference between the number of cells in G2M/S and G1 states in the liver and femur.

## KEY RESOURCES TABLE

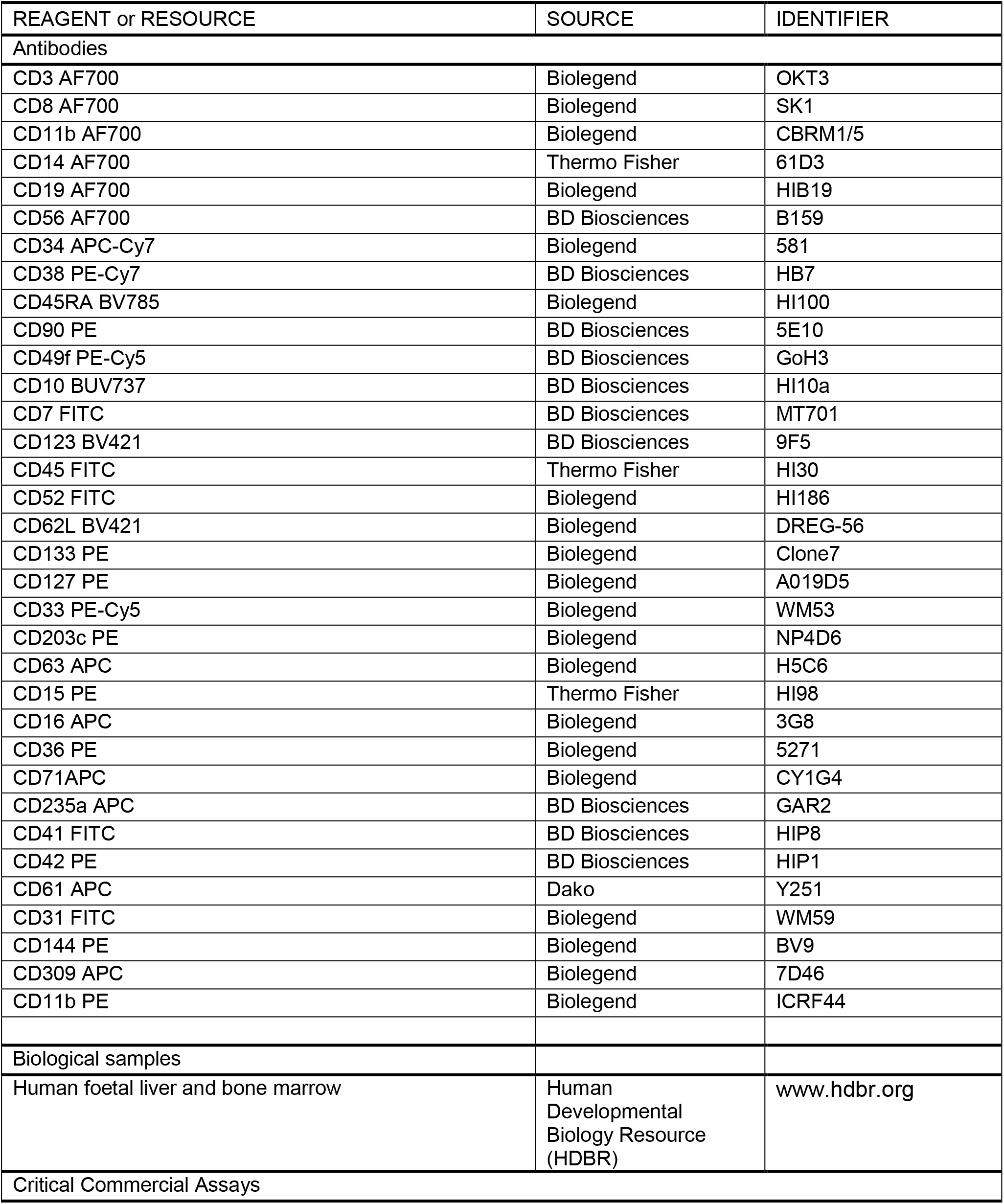
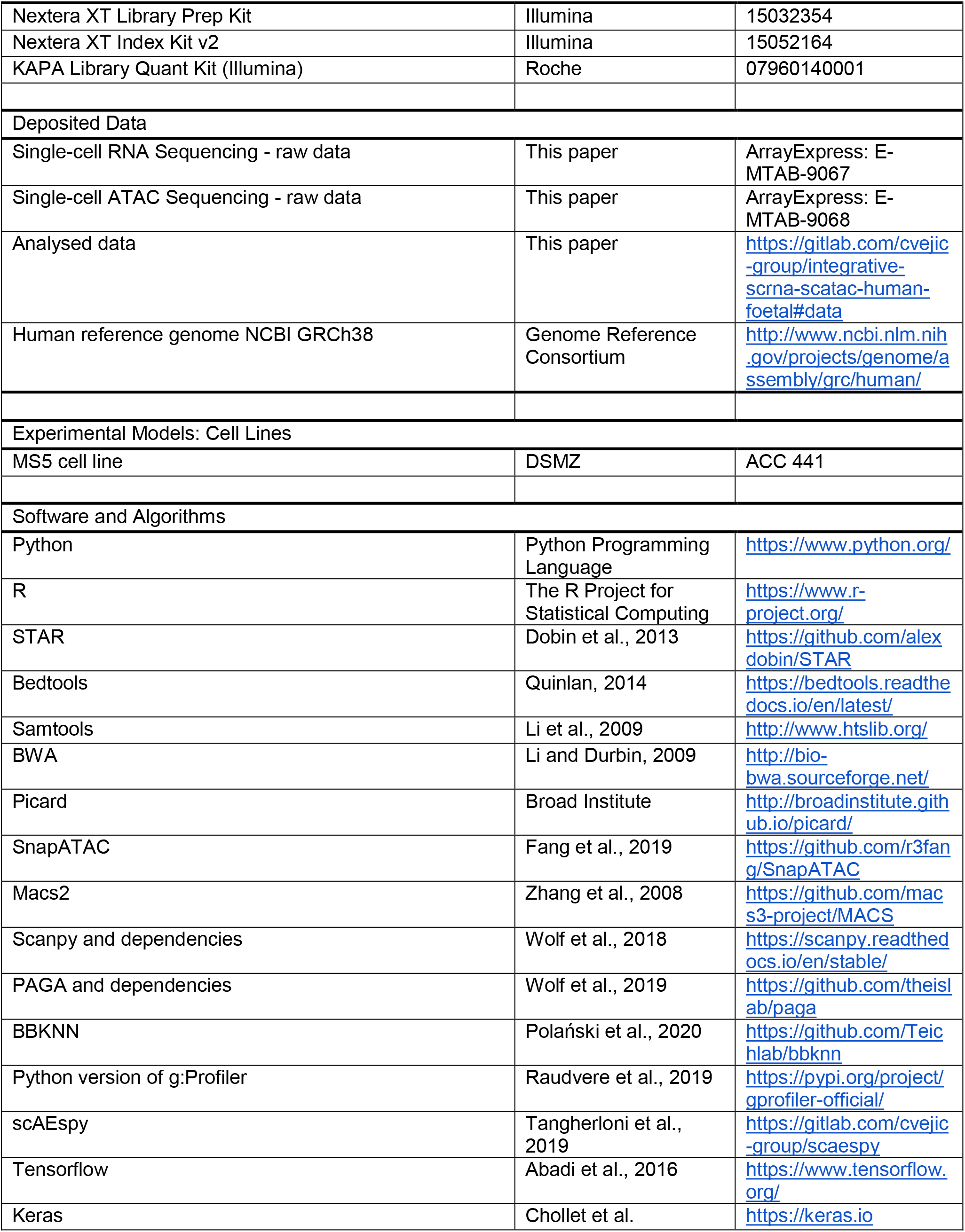
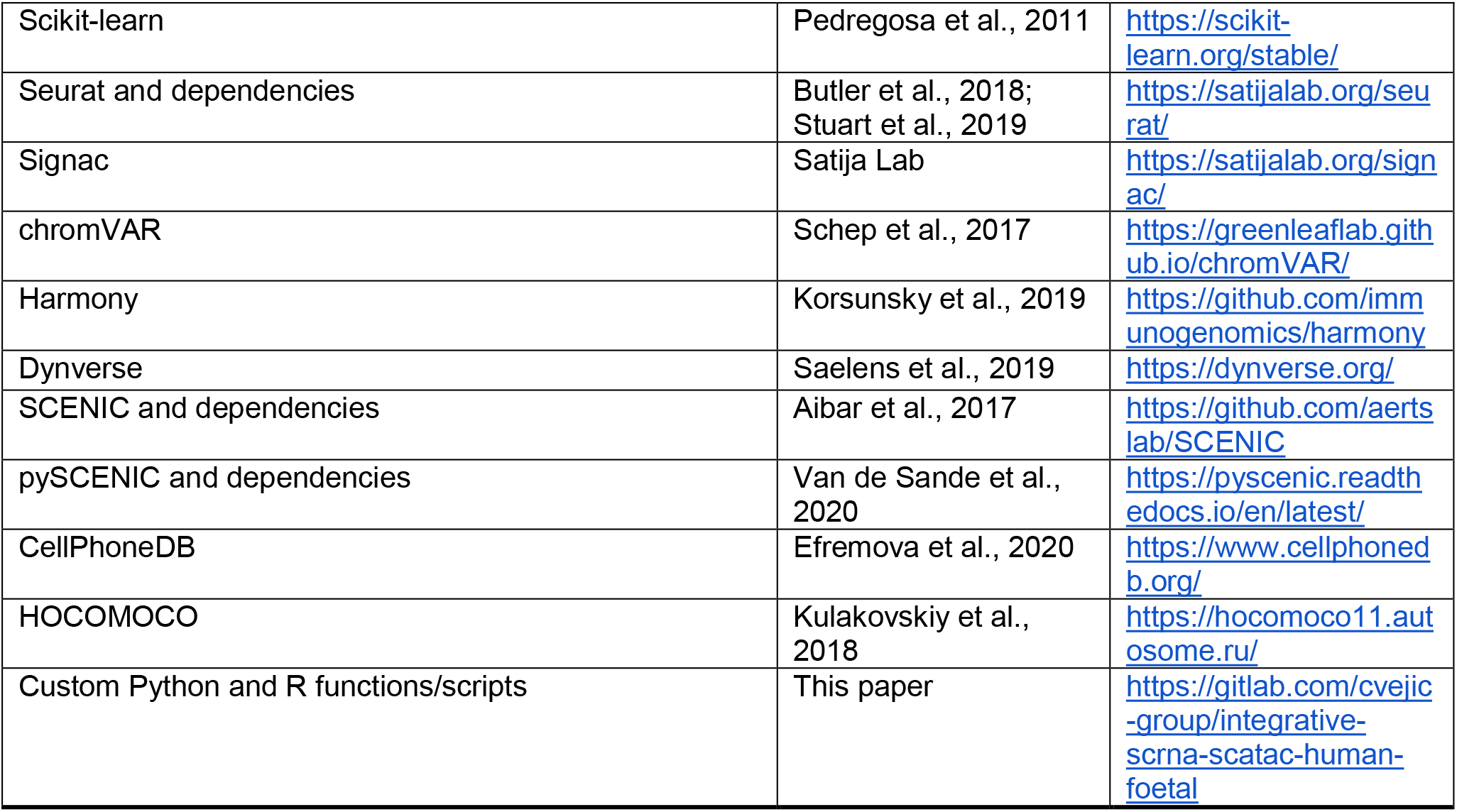

**Supplementary figure 1.**
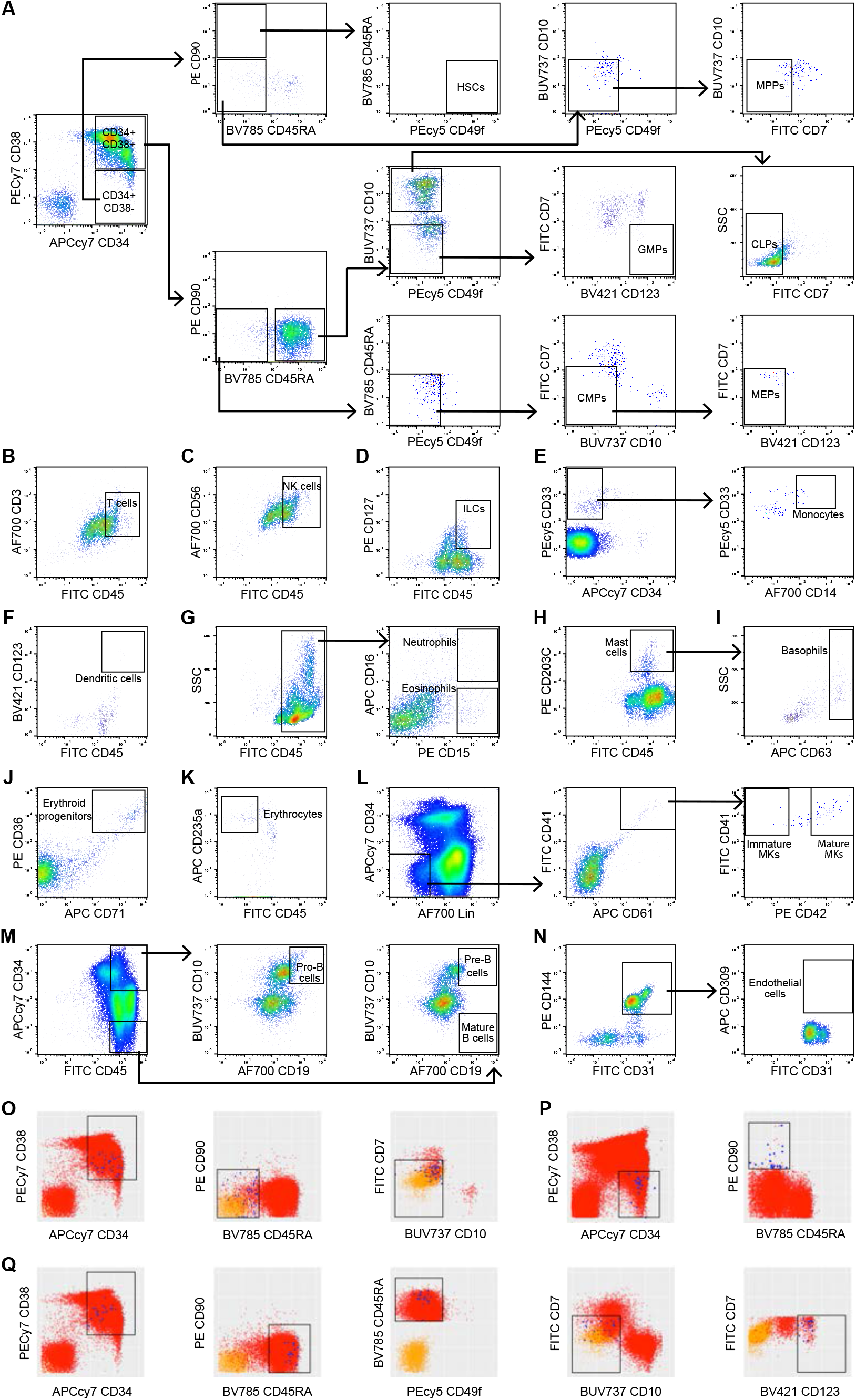
Sorting panels, Related to Figure 1, related to Figure 1. **A-N.** FACS sorting panel and gating strategy for the isolation of phenotypically defined cell types: **A.** Committed and non-committed haematopoietic progenitors, HSCs, MPPs, CMPs, GMPs, MEPs, and CLPs. **B.** T cells. **C.** NK cells. **D.** ILCs. **E.** Monocytes. **F.** Dendritic cells. **G.** Neutrophils and eosinophils. **H.** Mast cells. **I.** Basophils. **J.** Erythroid progenitors. **K.** Erythrocytes. **L.** Immature and mature MKs. **M.** Pro-B cells, pre-B cells, and mature B cells. **N.** Endothelial cells. **O-Q.** Index sorting data FACS plots showing sorted cells (blue), total gated population (red), and unstained population (yellow) for defined cell types: **O.** CMPs. **P.** HSCs. **Q.** GMPs. HSCs - haematopoietic stem cells, MPPs - multipotent progenitors, CMPs - common myeloid progenitors, GMPs - granulocyte-monocyte progenitors, MEPs - megakaryocyte-erythroid progenitors and CLPs - common lymphoid progenitors, NK cells - natural killer cells, ILCs - innate lymphoid cells, MKs - megakaryocytes.

**Supplementary figure 2.**
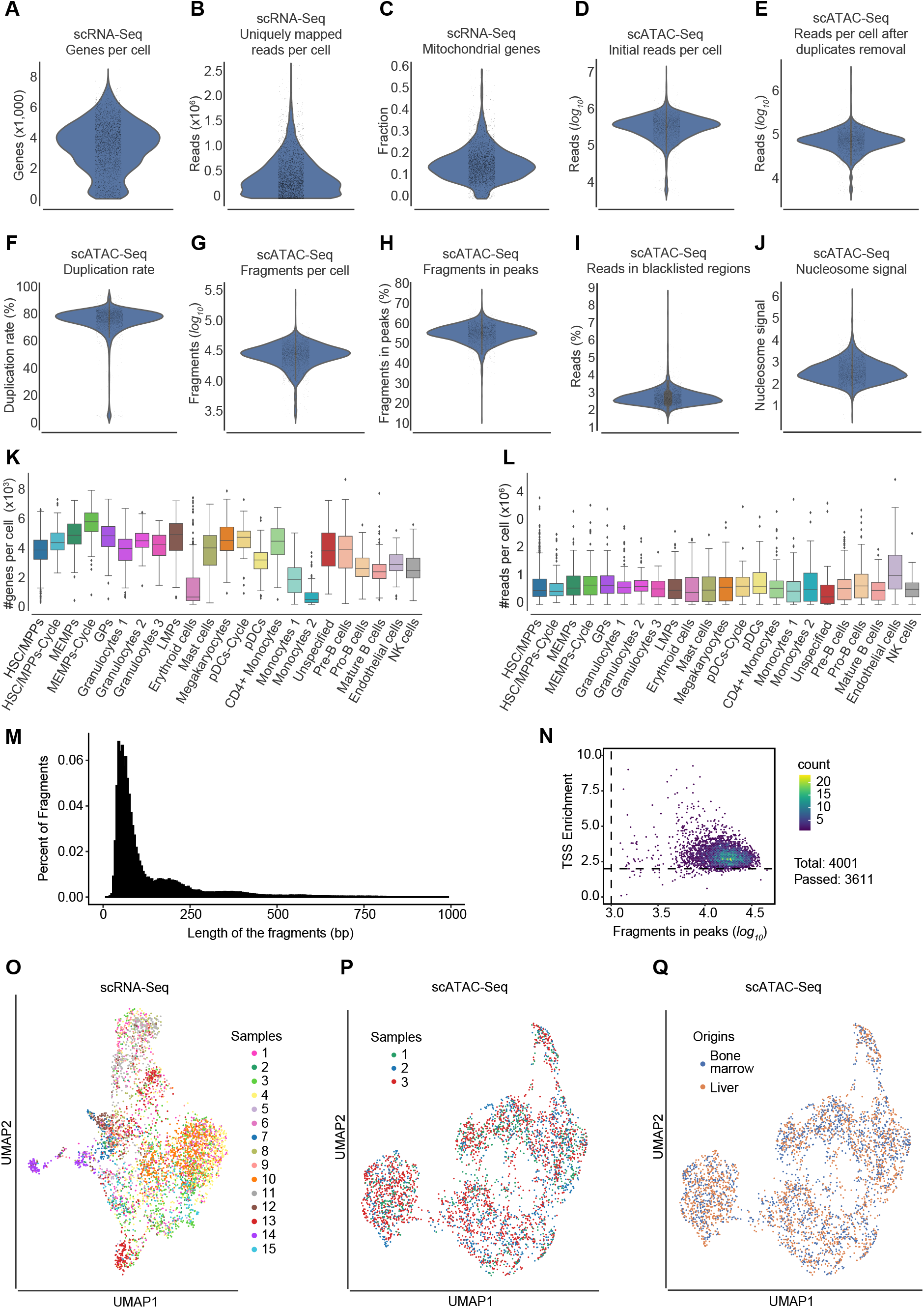
Quality control and batch effects correction in scRNA-Seq and scATAC-Seq data, related to Figures 1 and 3. **A.** Violin plots showing the number of expressed genes per cell in scRNA-Seq data. **B.** Violin plots showing the number of uniquely mapped reads against the reference genome per cell in scRNA-Seq data. **C.** Violin plots showing the fraction of mitochondrial genes compared to all genes per cell in scRNA-Seq data. **D.** Violin plots showing the number of reads per cell, prior to duplicates removal, in scATAC-Seq data. The y-axis is in *log*_10_ scale. **E.** Violin plots showing the number of reads per cell after duplicates removal in scATAC-Seq data. The y-axis is in *log*_10_ scale. **F.** Violin plots showing the duplicate rate in scATAC-Seq data. **G.** Violin plots showing the number of fragments per cell in scATAC-Seq data. The y-axis is in *log*_10_ scale. **H.** Violin plots showing the percentage of fragments per peak in scATAC-Seq data. **I.** Violin plots showing the percentage of reads mapping to the blacklist regions in scATAC-Seq data. **J.** Violin plots showing the nucleosome signal per cell in scATAC-Seq data. **K.** Box plot showing the number of genes per cell in each identified cell type in scRNA-Seq data. **L.** Box plot showing the number of uniquely mapped reads per cell in each identified cell type in scRNA-Seq data. M. Histogram showing the length of the fragments in terms of base pairs (200 bins). **N.** Scatterplot showing the fragments in peaks with respect to TSS enrichment. The colour intensity represents the number of counts. The x-axis is in *log*_10_ scale. **O.** UMAP visualization of the scRNA-Seq samples (*n* = 15) after the batch effect correction with BBKNN. Each colour represents a different sample. **P.** UMAP visualization of the scATAC-Seq samples (*n* = 3) after the batch effect correction with Harmony. Each colour represents a different sample. **Q.** UMAP visualization of the scATAC-Seq bone marrow (blue) and liver (orange) CD34+ CD38− cells after the batch effect correction with Harmony.

**Supplementary figure 3.**
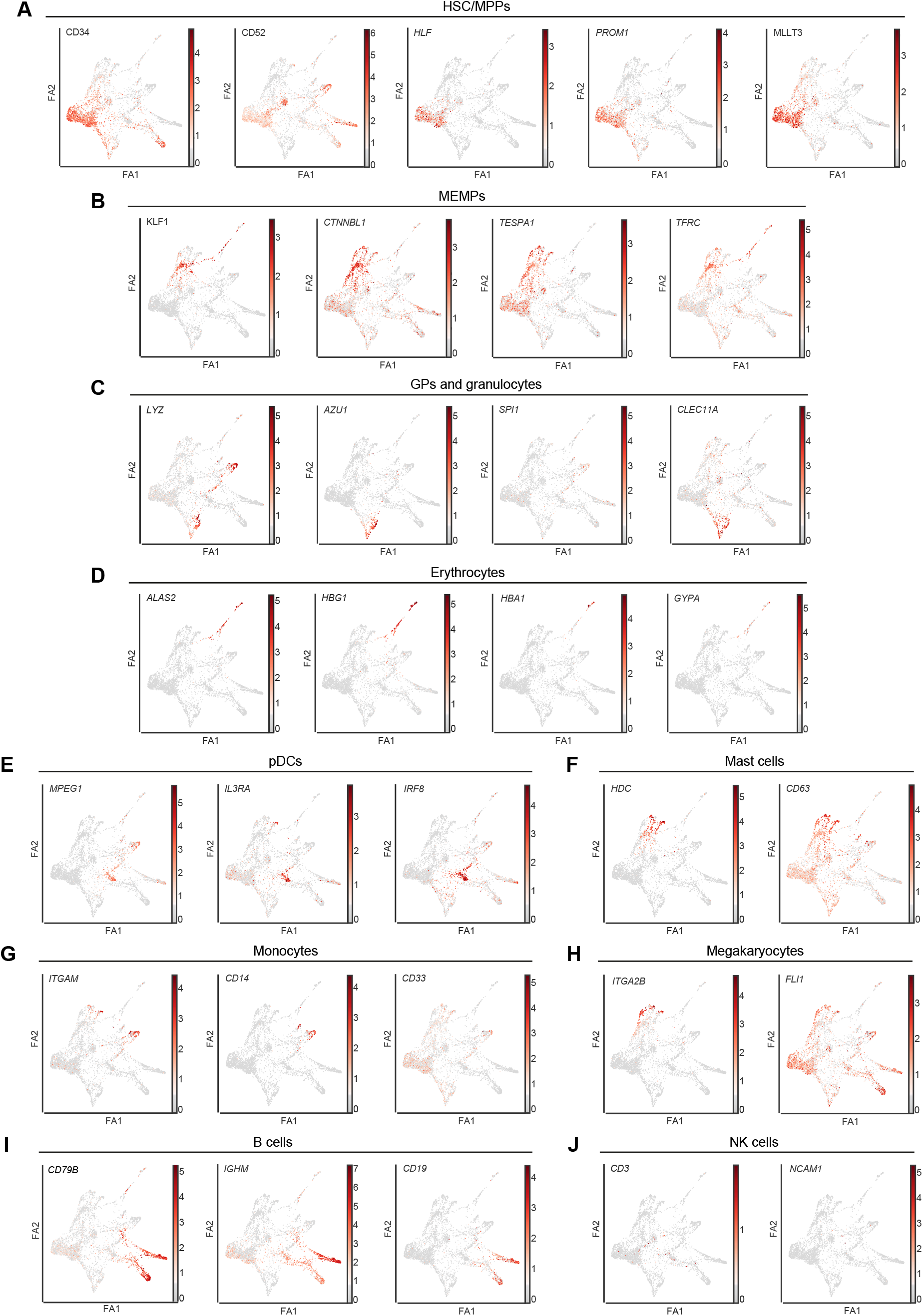
Expression of top marker genes along the differentiation trajectory, related to Figure 2. **(A-J)** FDG visualisation of the *log*-normalised gene expression of marker genes along the differentiation trajectory. **A.** HSC/MPPs (*CD34*, *CD52*, *HLF*, *PROM1*, and *MLLT3*). **B.** MEMPs (*KLF1*, *CTNNBL1*, *TESPA1*, and *TFRC*). **C.** GPs and granulocytes (*LYZ*, *AZU1*, *SPI1*, and *CLEC11A*). **D.** Erythrocytes (*ALAS2*, *HBG1*, *HBA1*, and *GYPA*). **E.** pDCs (*MPEG1*, *IL3RA*, and *IRF8*). **F.** Mast cells (*HDC* and *CD63*). **G.** Monocytes (*ITGAM*, *CD14*, and *CD33*). **H.** Megakaryocytes (*ITGA2B* and *FLI1*). **I.** B cells (*CD79B*, *IGHM*, and *CD19*). **J.** NK cells (*CD3* and *NCAM1*). Force-Directed Graph - FDG; ForceAtlas2 - FA2; HSC/MPPs - haematopoietic stem cells/multipotent progenitors; MEMPs - megakaryocyte-erythroid-mast progenitors; GPs - granulocytic progenitors; pDCs-Cycle - cycling plasmacytoid dendritic cells.

**Supplementary figure 4.**
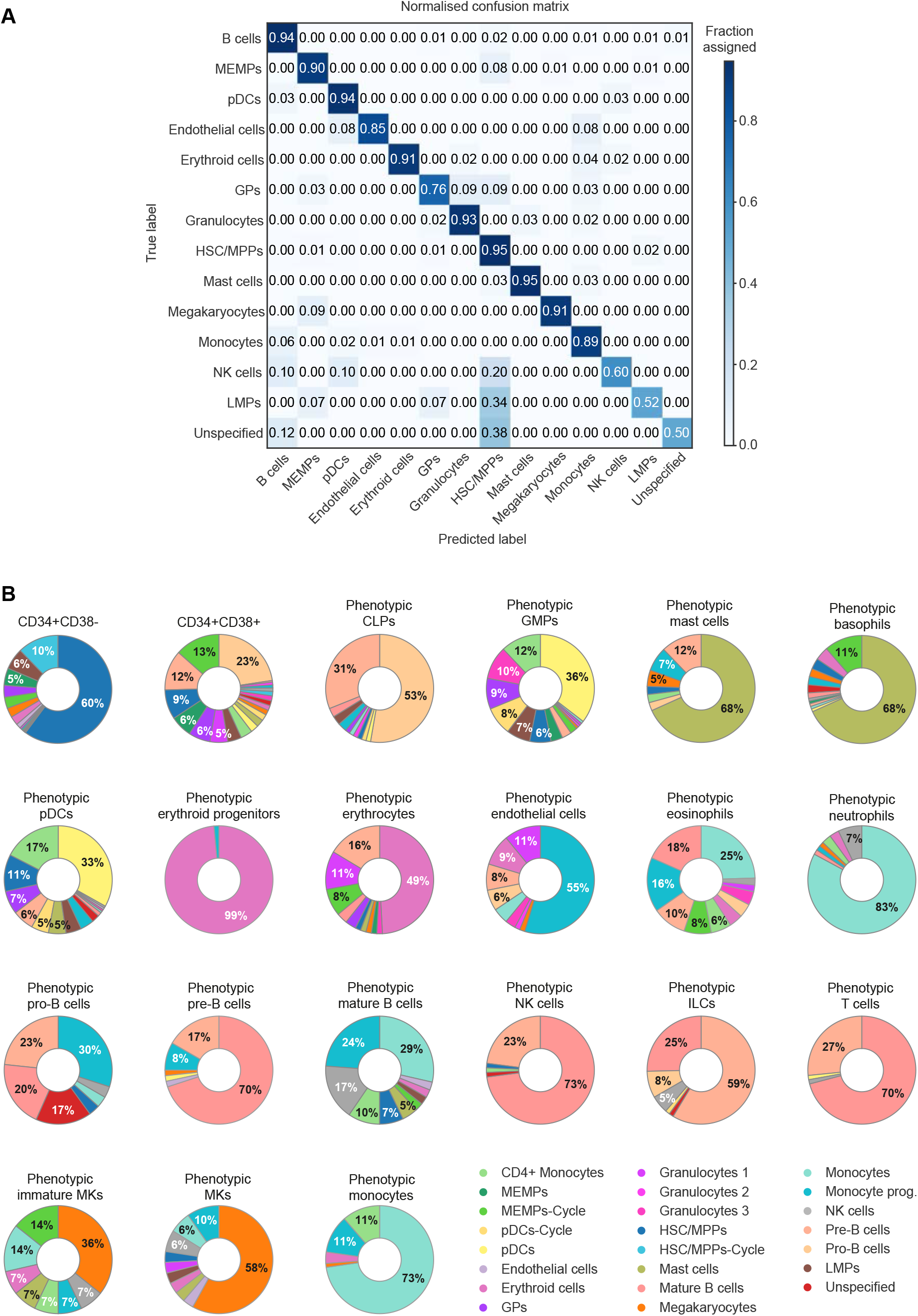
Validation of the cell type assignment and transcriptional heterogeneity of phenotypically defined cell populations, related to Figures 1 and 3. **A.** Confusion matrix showing the cell type assignment achieved by the DNN on the test set (901 cells), considering the top 30 marker genes per cell type (14 distinct cell types). The colour intensity represents the fraction of the assigned cells per cell type. **B.** Donut plots showing the percentage of transcriptionally defined (i.e., manually curated) cell populations in each of the phenotypically defined populations (Expanded from Figure 1C). Each colour represents a different cell type. HSC/MPPs-Cycle - cycling haematopoietic stem cells/multipotent progenitors; HSC/MPPs - haematopoietic stem cells/multipotent progenitors; MEMPs - megakaryocyte-erythroid-mast progenitors; MEMPs-Cycle - cycling megakaryocyte-erythroid-mast progenitors; GPs - granulocytic progenitors; LMPs - lymphomyeloid progenitors; pDCs-Cycle - cycling plasmacytoid dendritic cells; pDCs - plasmacytoid dendritic cells.

**Supplementary figure 5.**
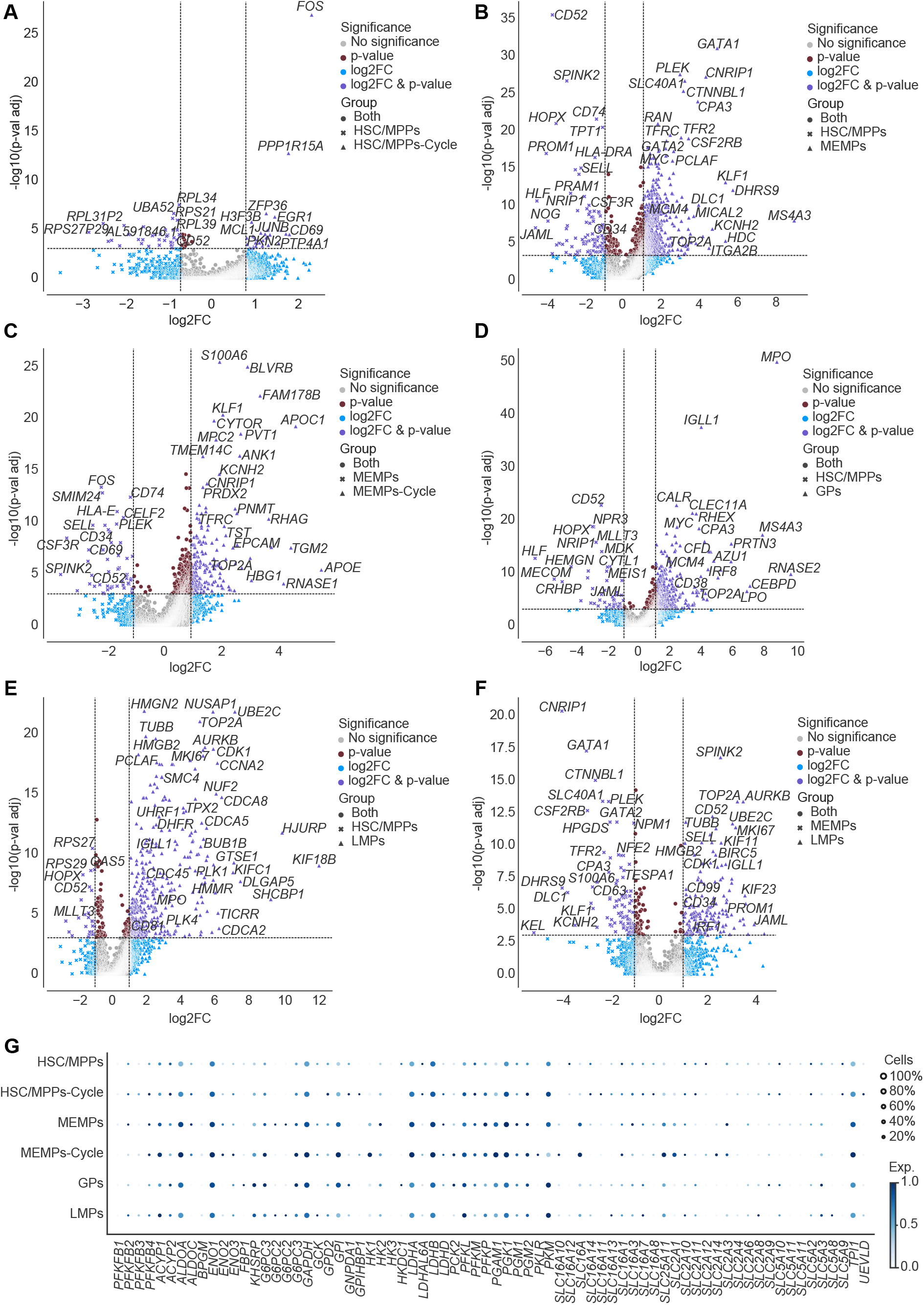
Differential expression analysis of the progenitor compartment, related to the STAR Methods section. **(A-F)** Volcano plot showing DEGs between two cell types of interest. **A.** HSC/MPPs and HSC/MPPs-Cycle. **B.** HSC/MPPs and MEMPs. **C.** MEMPs and MEMPs-Cycle. **D.** HSC/MPPs and GPs. **E.** HSC/MPPs and LMPs. **F.** MEMPs and LMPs. The x-axes show the *log*_2_ fold-change (magnitude of change), while the y-axes show the *log*_10_ adjusted p-value (statistical significance). We used the Wilcoxon rank-sum with the Benjamini-Hochberg correction. Colours represent the significance of the genes, both in terms of p-value and *log*_2_ fold-change. **G.** Dot plot of the expression of metabolic genes involved in glycolysis in the identified progenitor compartment. The expression of the genes is standardised between 0 and 1. For each gene, the minimum value is subtracted and the result is divided by the maximum. The spot size indicates the percentage of cells that express the gene of interest within each cell type. The colour intensity represents the standardised expression level. HSC/MPPs-Cycle - cycling haematopoietic stem cells/multipotent progenitors; HSC/MPPs - haematopoietic stem cells/multipotent progenitors; MEMPs - megakaryocyte-erythroid-mast progenitors; MEMPs-Cycle - cycling megakaryocyteerythroid-mast progenitors; GPs - granulocytic progenitors; LMPs - lympho-myeloid progenitors.

**Supplementary figure 6.**
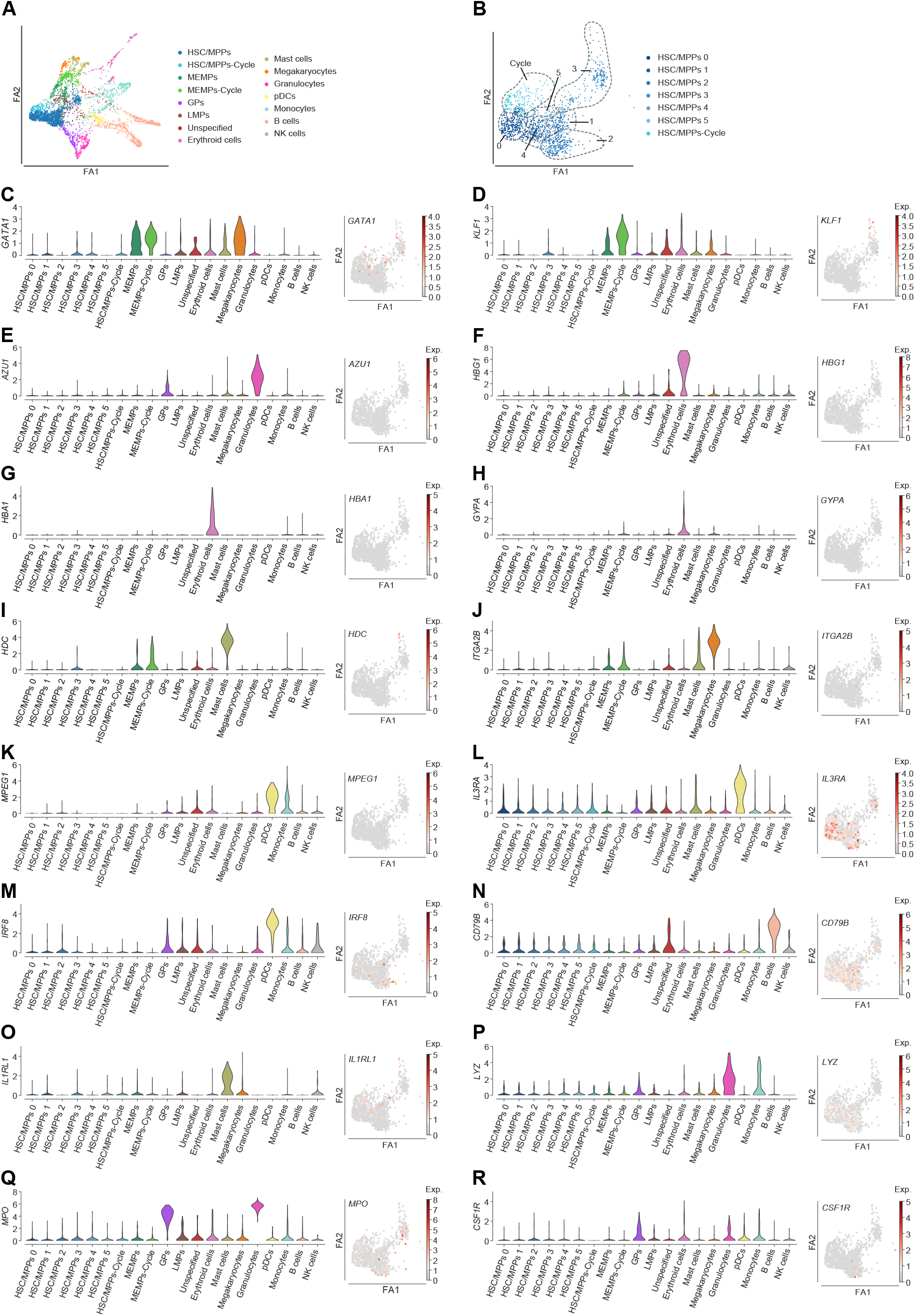
Expression of lineage-specific marker genes in HSC/MPP sub-populations, related to 2. **A.** FDG visualisation of the identified differentiation trajectory. **B.** FDG visualisation of the HSC/MPP sub-populations. **(C-R)** Left panels - violin plots showing the log-normalised median gene expression of lineage-specific marker genes in the HSC/MPP sub-populations and more differentiated haematopoietic cells. Right panels - FDG visualisations of the log-normalised gene expression of the same genes as in violin plots along the differentiation trajectory, considering only the HSC/MPP sub-populations. **C.** *GATA1*. **D.** *KLF1*. **E.** *AZU1*. **F.** *HBG1*. **G.** *HBA1*. **H.** *GYPA*. **I.** *HDC*. **J.** *ITGA2B*. **K.** *MPEG1*. **L.** *IL3RA*. **M.** *IRF8*. **N.** *CD79B*. **O.** *IL1RL1*. **P.** *LYZ*. **Q.** *MPO*. **R.** *CSF1R*. Force-Directed Graph - FDG; ForceAtlas2 - FA2; HSC/MPPs - haematopoietic stem cells/multipotent progenitors; HSC/MPPs-Cycle - cycling haematopoietic stem cells/multipotent progenitors; MEMPs - megakaryocyte-erythroid-mast progenitors; MEMPs-Cycle - cycling megakaryocyte-erythroid-mast progenitors; GPs - granulocytic progenitors; LMPs - lymphomyeloid progenitors; pDCs - plasmacytoid dendritic cells.

**Supplementary figure 7.**
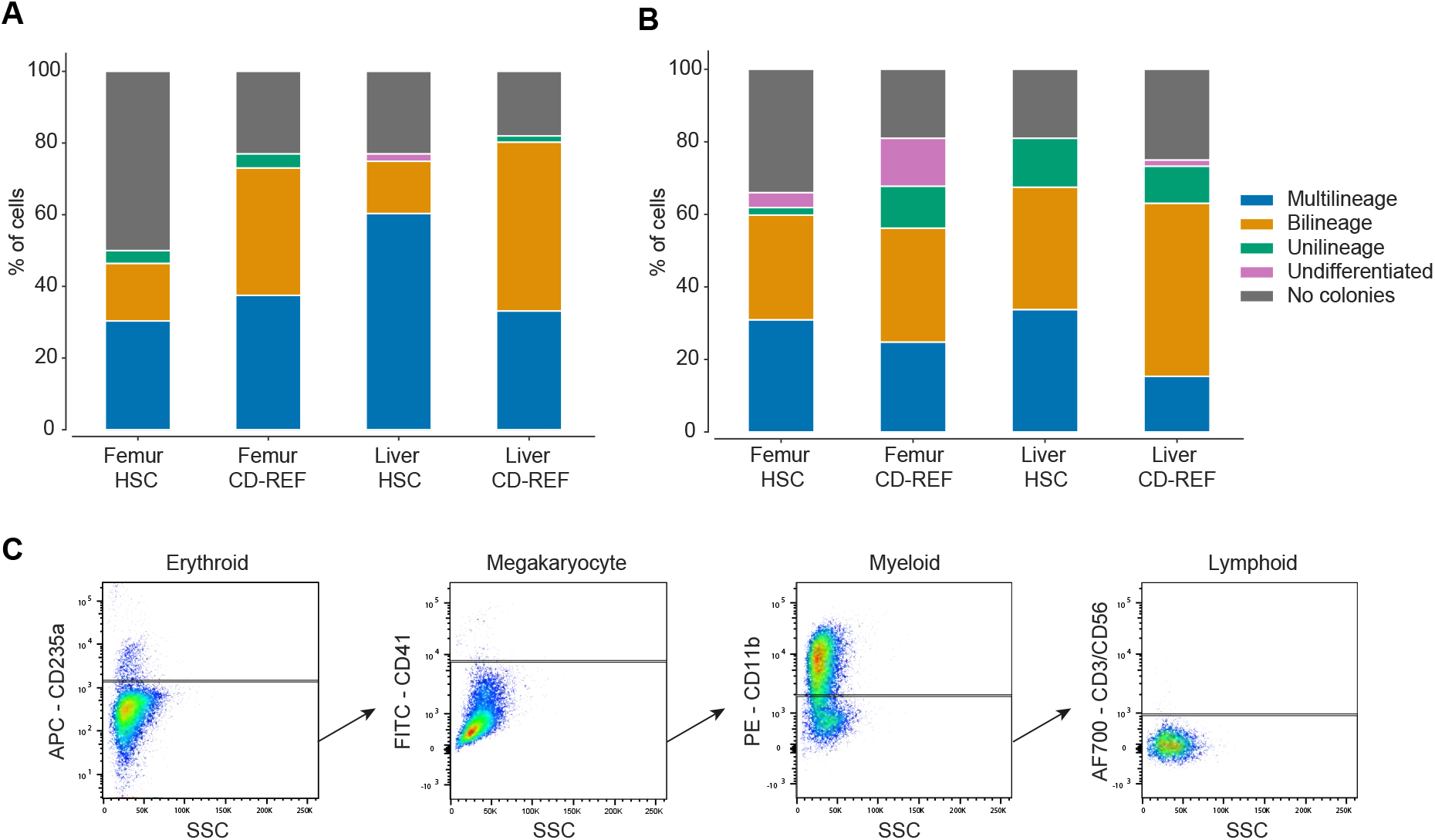
Colony formation and the lineage output of CD-REF cells and phenotypic HSCs isolated from the foetal liver and femur, related to Figure 6. **A-B.** Stacked bar chart showing the differentiation potential and efficiency of colony formation of CD-REF cells compared to the phenotypic HSCs isolated from foetal liver and femur. The y-axis shows the percentage of colonies. The colonies have been divided by their differentiation potential as determined by FACS, namely: multilineage, bilineage, unilineage, and undifferentiated. Data from two different experiments are shown. **C.** Representative FACS plot showing the lineage composition of a quadrilineage colony.

**Supplementary table 1.** Cell-surface markers used to isolate cell types, related to Figure 1.

| Phenotypic cell type | Cell-surface markers |
| --- | --- |
| HSCs | Lin- CD34+ CD38- CD45RA- CD90+ CD49f+/- |
| MPPs | Lin- CD34+ CD38- CD90- CD45RA- CD49f- CD10- CD7- |
| CMPs | Lin- CD34+ CD38+ CD90- CD45RA- CD49f- CD10- CD7- |
| MEPs | Lin- CD34+ CD38+ CD90- CD45RA- CD49f- CD10- CD7- CD123- |
| GMPs | Lin- CD34+ CD38+ CD90- CD45RA+ CD49f- CD10- CD7- CD123+/- |
| CLPs | Lin-, CD34+ CD38+ CD90- CD45RA+ CD49f - CD10+ CD7- |
| T cells | CD45+ CD3+ |
| NK cells | CD45+ CD56+ |
| ILCs | Lin+/- CD45+ CD127+ |
| Monocytes | CD34- CD33+ CD14+ |
| Dendritic cells | Lin- CD45+ CD123+ |
| Mast cells | CD45+ CD203c+ |
| Basophils | CD45+ CD203c+ CD63+ |
| Neutrophils | CD45+ CD15+ CD16+ |
| Eosinophils | CD45+ CD15+ CD16- |
| Erythroid progenitors | CD36+ CD71+ |
| Erythrocytes | CD235a+ |
| Immature MKs | Lin- CD34- CD41+ CD61+ CD42- |
| Mature MKs | Lin- CD34- CD41+ CD61+ CD42+ |
| Pro-B cells | CD45+ CD34+ CD19+ CD10+ |
| Pre-B cells | CD45+ CD34- CD19+ CD10+ |
| Mature B cells | CD45+ CD34- CD19+ CD10- |
| Endothelial cells | CD31+ CD144+ CD309+ |

**Supplementary table 2.**
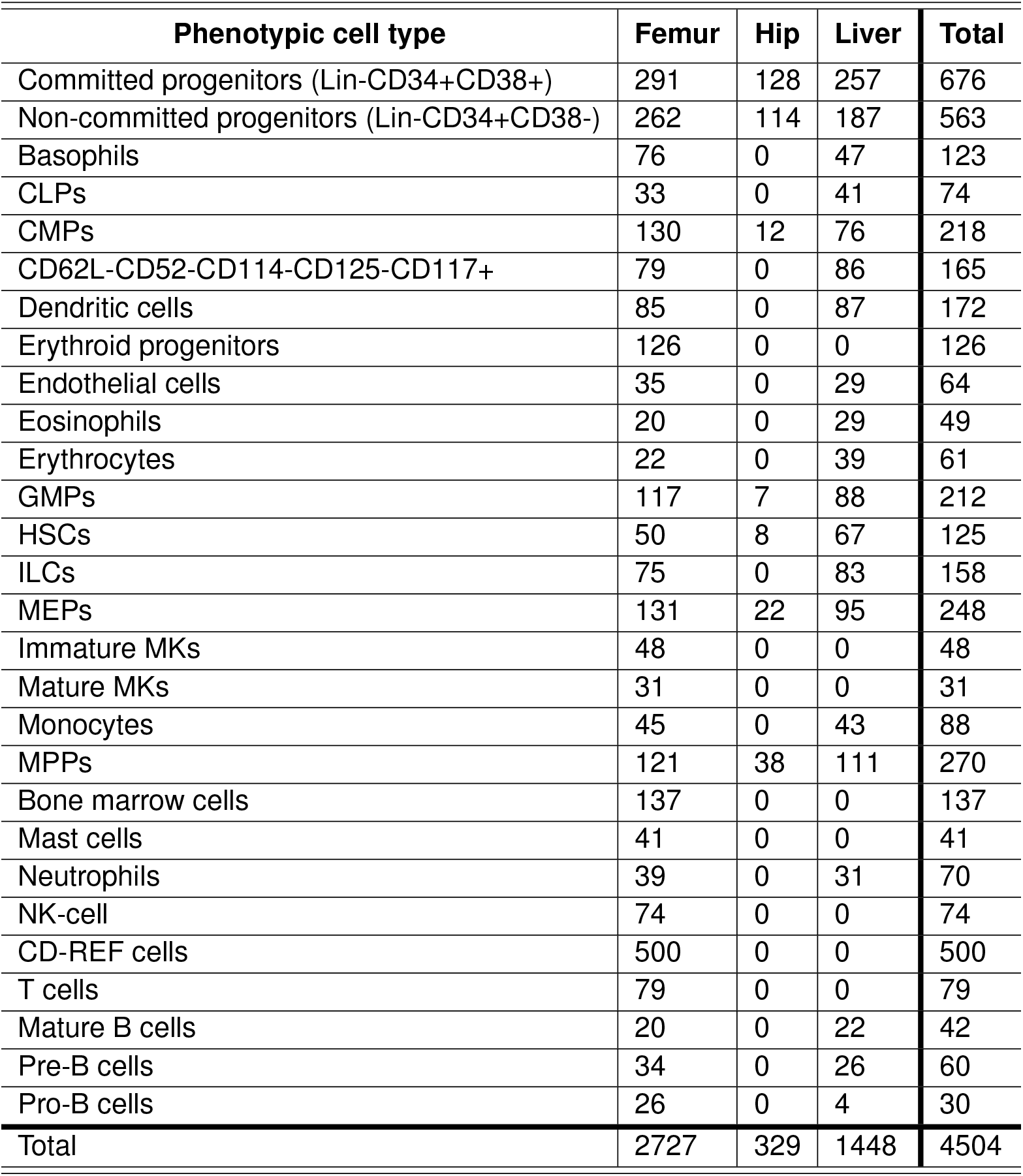
Phenotypic cell types per organ, related to Figure 1.

| **Phenotypic cell type** | **Femur** | **Hip** | **Liver** | **Total** |
| --- | --- | --- | --- | --- |
| Committed progenitors (Lin-CD34+CD38+) | 291 | 128 | 257 | 676 |
| Non-committed progenitors (Lin-CD34+CD38-) | 262 | 114 | 187 | 563 |
| Basophils | 76 | 0 | 47 | 123 |
| CLPs | 33 | 0 | 41 | 74 |
| CMPs | 130 | 12 | 76 | 218 |
| CD62L-CD52-CD114-CD125-CD117+ | 79 | 0 | 86 | 165 |
| Dendritic cells | 85 | 0 | 87 | 172 |
| Erythroid progenitors | 126 | 0 | 0 | 126 |
| Endothelial cells | 35 | 0 | 29 | 64 |
| Eosinophils | 20 | 0 | 29 | 49 |
| Erythrocytes | 22 | 0 | 39 | 61 |
| GMPs | 117 | 7 | 88 | 212 |
| HSCs | 50 | 8 | 67 | 125 |
| ILCs | 75 | 0 | 83 | 158 |
| MEPs | 131 | 22 | 95 | 248 |
| Immature MKs | 48 | 0 | 0 | 48 |
| Mature MKs | 31 | 0 | 0 | 31 |
| Monocytes | 45 | 0 | 43 | 88 |
| MPPs | 121 | 38 | 111 | 270 |
| Bone marrow cells | 137 | 0 | 0 | 137 |
| Mast cells | 41 | 0 | 0 | 41 |
| Neutrophils | 39 | 0 | 31 | 70 |
| NK-cell | 74 | 0 | 0 | 74 |
| CD-REF cells | 500 | 0 | 0 | 500 |
| T cells | 79 | 0 | 0 | 79 |
| Mature B cells | 20 | 0 | 22 | 42 |
| Pre-B cells | 34 | 0 | 26 | 60 |
| Pro-B cells | 26 | 0 | 4 | 30 |
| **Total** | **2727** | **329** | **1448** | **4504** |

**Supplementary table 3.** scRNA-Seq samples, related to the STAR Methods section.

| **No.** | **Gates** | **Min**<br>**#counts** | **Max**<br>**#counts** | **Min**<br>**#genes** | **Max**<br>**#genes** | **Max**<br>**%mito** |
| --- | --- | --- | --- | --- | --- | --- |
| 1 | Non-committed progenitors<br>Committed progenitors | 10,000 | 1,750,000 | 200 | 7,500 | 40 |
| 2 | Non-committed progenitors<br>Committed progenitors<br>HSCs | 10,000 | 2,000,000 | 200 | 7,500 | 40 |
| 3 | Non-committed progenitors<br>Committed progenitors<br>HSCs<br>MPPs<br>CMPs<br>GMPs<br>MEPs | 10,000 | 1,500,000 | 200 | 7,500 | 40 |
| 4 | Non-committed progenitors<br>Committed progenitors<br>HSCs<br>MPPs<br>CMPs<br>GMPs<br>MEPs | 10,000 | 2,750,000 | 200 | 9,000 | 40 |
| 5 | Immature MKs<br>Mature MKs | 10,000 | 1,750,000 | 200 | 9,000 | 40 |
| 6 | Non-committed progenitors<br>Committed progenitors<br>HSCs<br>MPPs<br>CMPs<br>GMPs<br>MEPs | 10,000 | 1,500,000 | 350 | 7,500 | 40 |
| 7 | Bone marrow cells | 400 | 1,750,000 | 200 | 7,500 | 60 |
| 8 | Endothelial cells<br>Pro-B cells<br>Pre-B cells<br>Mature B | 400 | 1,500,000 | 200 | 7,500 | 60 |
| 9 | CLPs<br>GMPs<br>Monocytes | 1,000 | 2,000,000 | 200 | 7,500 | 40 |
| 10 | CD-REF cells | 1,000 | 550,000 | 200 | 6,000 | 40 |
| 11 | ILCs<br>NK cells<br>T cells | 2,000 | 1,500,000 | 200 | 10,000 | 40 |
| 12 | Neutrophils<br>Eosinophils<br>Erythrocytes | 2,000 | 1,500,000 | 200 | 6,000 | 40 |
| 13 | Dendritic cells<br>Mast cells<br>Basophils | 2,000 | 1,500,000 | 200 | 7,500 | 40 |
| 14 | Erythroid progenitors | 500 | 3,000,000 | 100 | 4,500 | 40 |
| 15 | CD62L- CD52- CD114-<br>CD125- CD117+ | 2,000 | 1,000,000 | 1,200 | 7,000 | 40 |

## Footnotes

1 Chollet *et al.*: https://keras.io

2 Broad Institute: http://broadinstitute.github.io/picard/

3 Stuart *et al.*: https://github.com/timoast/signac/

